# Widespread reorganisation of the regulatory chromatin landscape facilitates resistance to inhibition of oncogenic ERBB2 signalling

**DOI:** 10.1101/2021.04.29.441944

**Authors:** Samuel Ogden, Kashmala Carys, Jason Bruce, the OCCAMS consortium, Andrew D. Sharrocks

**Affiliations:** School of Biological Sciences, Faculty of Biology, Medicine and Health, University of Manchester, Michael Smith Building, Oxford Road, Manchester, M13 9PT, UK

**Author notes:** Corresponding author: Andrew Sharrocks.

**Keywords:** Oesophageal adenocarcinoma, chromatin, ERBB2, PPARGC1A, HNF4A, drug resistance

## Abstract

Oesophageal adenocarcinoma (OAC) patients show poor survival rates and there are few targeted molecular therapies available. However, components of the receptor tyrosine kinase (RTK) driven pathways are commonly mutated in OAC, typified by high frequency amplifications of the RTK *ERRB2*. ERBB2 can be therapeutically targeted, but this has limited clinical benefit due to the acquisition of drug resistance. Here we examined how OAC cells respond to ERBB2 inhibition through altering their regulatory chromatin landscapes and rewiring their gene regulatory networks to acquire a reversible resistant state. ERBB2 inhibition triggers widespread remodelling of the accessible chromatin landscape. This remodelling is accompanied by the activation of the transcriptional regulators HNF4A and PPARGC1A. Initially, inhibition of cell cycle associated gene expression programmes is observed, with compensatory increases in the programmes driving changes in metabolic activity. PPARGC1A is instrumental in promoting a switch to dependency on oxidative phosphorylation and both PPARGC1A and HNF4A are required for the acquisition of resistance to ERBB2 inhibition. Our work therefore reveals the molecular pathways that support the acquisition of a resistant state and points to potential new therapeutic strategies to combat drug resistance.

## Introduction

Cancer is predominantly caused by DNA mutations and genomic rearrangements. However, it is becoming increasingly clear that rewiring of the epigenetic landscape also plays a pivotal role in tumourigenesis (Somerville et al., 2018, Flavahan et al., 2019; reviewed in Allis and Jenuwein, 2016; Jones et al., 2016). This epigenetic rewiring leads to changes in gene expression programmes and the molecular pathways that are operational in the cell (reviewed in Bradner et al., 2017). The molecular changes manifested by a cancer cell provide the opportunity for personalised treatment which is exemplified by the use of inhibitors like trastuzumab and lapatinib to treat cancer patients with amplifications of the RTK ERBB2 (Blackwell et al., 2010; Cobleigh et al., 1999). However, drug resistance often arises, limiting the effectiveness of treatment. This is especially the case for OAC where trastuzumab and lapatinib have been shown to have relatively limited effects on patient survival (Bang et al., 2010; Hecht et al., 2016). Drug resistance in OAC often arises due to the selection of cells containing compensatory mutations (Kim et al., 2014; Janjigian et al., 2018; Wang et al., 2019). However, it is becoming increasingly recognised that changes to the epigenetic landscape can also play an important role in drug resistance, particularly in enabling the survival of “persistor” cells, which ultimately gather additional mutations to adopt a stable resistant form (Bi et al., 2020; Sharma et al., 2010; Ramirez et al., 2016; reviewed in Hammerlindl and Schaider, 2018; Shen et al., 2020).

The incidence of OAC is rapidly expanding in the Western world while the survival rates remain poor (Coleman et al., 2018). A pre-malignant state known as Barrett’s oesophagus (BO) is thought to be the precursor to OAC (Peters et al., 2019). Genome sequencing studies have revealed numerous mutational changes in the transition from BO to OAC but there are few high frequency recurrent oncogenic driver events (Weaver et al., 2014; Stachler et al., 2018; Frankell et al., 2019). However, at the pathway level, components of the receptor tyrosine kinase (RTK) driven pathways are frequently mutated in OAC (60%-76%; Cancer Genome Atlas Research Network et al, 2017; Frankell et al., 2019), typified by relatively high frequency amplifications and mutations of the RTK ERRB2 (ranging from 18-32% tumours; Cancer Genome Atlas Research Network et al., 2017; Frankell et al., 2019). At the epigenetic level, the accessible chromatin landscape of BO is vastly different to the surrounding normal oesophageal tissue and this landscape is further altered during the transition to OAC (Rogerson et al., 2019; 2020). These chromatin changes are accompanied by alterations to the transcriptional regulatory networks, with many changes to transcription factor activity being common to BO and OAC, including transcription factors like HNF4A, GATA6, FOXA, HNF1B and PPARG (Rogerson et al., 2019; Chen et al., 2020; Ma et al., 2021). Conversely, other transcription factors appear more dominant in OAC such as AP1 (Britton et al., 2017) or their regulatory activity is repurposed and directed to alternative transcriptional programmes as exemplified by KLF5 (Rogerson et al., 2020). However, it is unclear how the deregulated RTK pathways impact on these gene regulatory networks.

Here we have investigated how inhibition of the RTK ERBB2 influences gene regulatory networks as OAC cells acquire drug resistance. We demonstrate rapid and widespread remodelling of the accessible chromatin landscape in response to ERBB2 inhibition. This revealed changes in transcriptional regulatory activities, which converged on HNF4A and PPARGC1A and their influence on metabolic programmes. Both HNF4A and PPARGC1A are required for the acquisition of a drug resistant state and their target gene networks represent potential therapeutic vulnerabilities to prevent the emergence of drug resistance.

## Results

### *ERRB2* amplified OAC cells develop resistance to lapatinib

Previous studies indicated that several gastro-oesophageal adenocarcinoma cancer cell lines harbour amplifications of the gene encoding ERBB2. By using ATAC-seq data, we validated this amplification event which encompasses the *ERRB2* locus in all of the cancer lines but is absent in the HET1A and CPA cell lines, derived from normal and Barrett’s oesophageal tissue respectively (Supplementary Fig. S1A). These amplifications lead to variable levels of ERBB2 protein expression with the highest levels in the OE19 and NCI-N87 cell lines (Fig. 1A). To select cell lines which most closely resembled patient derived samples, we analysed the open chromatin landscapes of the cell line panel compared to three OAC patient biopsies which harbour *ERBB2* amplifications. Principal component analysis demonstrated tight clustering of samples from normal or Barrett’s oesophagus, and a distinct looser cluster of the OAC samples (Fig. 1B). The OE19, ESO26, KYAE1 and OAC organoid WTSI-OESO_009 (CAM408, Li et al., 2018) cluster together with these tumour samples whereas OE33 and NCI-N87 are more distantly associated. Furthermore, clustering based on Pearson’s correlations of the same data gave the same broad conclusions with OE33 cells being a clear outlier. Importantly, we also examined the expression of a set of transcription factors we previously associated with Barrett’s and OAC (Rogerson et al., 2019) and two markers of squamous epithelium, *TP63* and *PAX9*, in a range of patient samples (Maag et al., 2017) and cell lines (Supplementary Fig. S1B). OE19, ESO26, and KYAE1 again clustered with OAC patient samples and exhibited expression of *GATA6*, *FOXA2* and *HNF4A*. We therefore took OE19, ESO26, and KYAE1 cells forward for further analysis as representative examples of *ERBB2*-amplified OAC.

**Figure 1.**
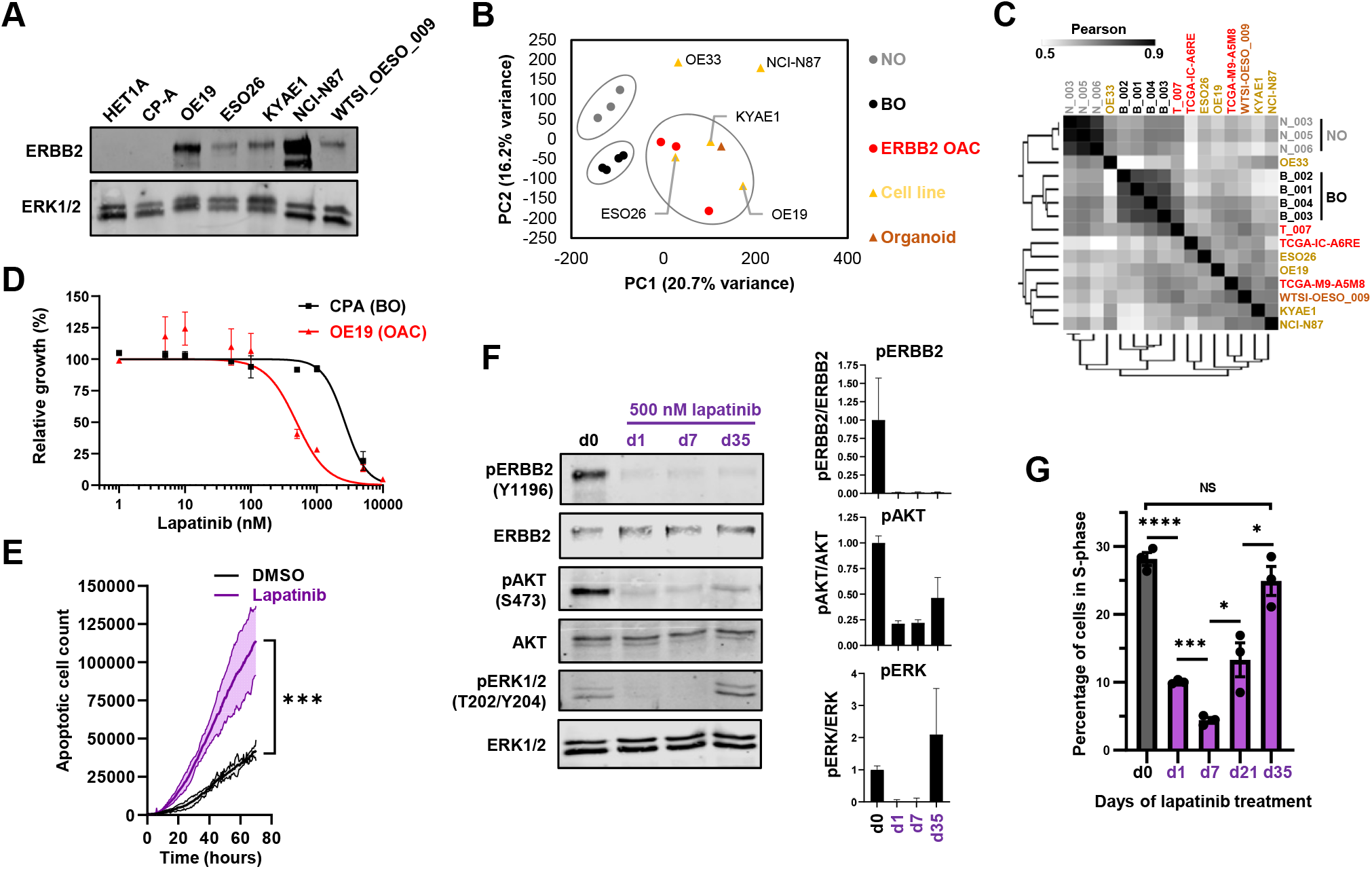
Resistance to ERBB2 inhibition in OAC cells arises after weeks of treatment. (A) Western blot for ERBB2 expression in the indicated cell lines. (B) Principal component analysis of human tissue and cell line ATAC-seq data. A union peakset composed of the top 50K peaks from BO and OAC tissue was used. NO – normal oesophageal tissue, BO – Barrett’s oesophageal tissue, ERBB2 OAC – oesophageal adenocarcinoma tissue harbouring *ERBB2* amplification, organoid – CAM408/WTSI-OESO_009. (C) Pearson correlation of ATAC-seq data shown in B. (D) MTS growth assay of OE19 cells treated with increasing concentrations of lapatinib for 72 hours. (E) Apoptosis assay of OE19 cells treated with vehicle control (DMSO) or 500 nM lapatinib for 70 hours. *** *P* < 0.001, 2-way ANOVA. 95% confidence intervals are shown, *n* = 3. (F) Western blot of OE19 cells treated with 500 nM lapatinib for the indicated timepoints: d0 – 24 hours DMSO, d1/7/35 – 1/7/35 days lapatinib. Quantification is shown, *n* = 3, error bars depict standard error of the mean (SEM). (G) Cell cycle analysis of OE19 cells treated with DMSO for 24 hours (d0) or 500 nM lapatinib for the indicated number of days. * *P* < 0.05, *** *P* < 0.001, **** *P* < 0.0001, unpaired t-test, error bars depict SEM, *n* = 3.

Next, we studied the response of OAC cells to treatment with the ERBB2/EGFR inhibitor lapatinib (Xia et al., 2002). In all cell lines, lapatinib caused a reduction in the activating phosphorylation (Y1196) of ERBB2 and downstream ERK1/2 (Supplementary Fig. S1C). In addition to signalling via ERK, ERBB2 also signals via a parallel pathway involving AKT and similar reductions in AKT phosphorylation levels were observed with the exception of ESO26 cells where co-treatment with an AKT inhibitor was required to gain maximal inhibition due to the presence of activating mutations in *PIK3CA* in this cell line (Kim et al., 2014). Genetic depletion of ERBB2 caused similar reductions in ERK and AKT phosphorylation in OE19 cells (Supplementary Fig. S1D). Lapatinib treatment caused dose-dependent reductions in growth of all the OAC lines (Fig. 1D; Supplementary Fig. S1E). Part of this reduced growth was due to increased levels of apoptosis as exemplified by OE19 cells (Fig. 1E). Having established an optimal concentration of lapatinib, we examined the response of OE19 cells over a 5 week period. ERBB2 phosphorylation remained inhibited for the duration of treatment but both AKT and ERK phosphorylation levels partially recovered after 5 weeks treatment (Fig. 1F). This recovery was accompanied by a resumption of proliferation as exemplified by the numbers of cells in S phase (Fig. 1G; Supplementary Fig. S1F). Thus by 5 weeks, the OE19 cells can be considered as resistant to ERBB2 inhibition with lapatinib and represent a good model for further investigation of the underlying resistance mechanisms.

### ERBB2 inhibition causes widespread changes to the transcriptional regulatory landscape in OAC cells

To understand how ERRB2 inhibition rewires OAC cells along the path to resistance, we focussed on OE19 cells and examined changes to their gene expression profile and associated open chromatin landscape following treatment with lapatinib for 1, 7 and 35 days (Fig. 2A). RNA-seq analysis revealed over a thousand genes changing expression (>2 fold change, FDR <0.05) after 1 day and 7 days (roughly equally split as up- and down-regulated), with a slightly lower number at 35 days (Fig. 2B; Supplementary Fig. 2A; Supplementary Table 1). Clustering based on PCA analysis and Pearson’s correlations indicated that by day 35, the transcriptome had begun to resemble the starting population (Supplementary Fig. S2B and C) which is also reflected in the recovery in expression levels of the differentially expressed genes (Fig. 2C and D; Supplementary Fig. S2D). Importantly, there is a substantial overlap in gene expression changes observed after 1 day of lapatinib treatment and RNAi-mediated ERBB2 depletion, which demonstrates the specificity of response to the inhibitor (Supplementary Fig. S2E). Gene ontology analysis of gene expression changes between sequential timepoints revealed a decrease in cell cycle gene expression which was further reduced at 7 days but reversed at 35 days as cells become resistant to ERBB2 inhibition (Fig. 2E) as exemplified by *FOXM1*, *MYBL2* and *MYC* expression (Fig. 2F). These genes are generally typical of cluster 5 (Fig. 2C and D). Conversely, different processes are upregulated at day 1, including metabolic and signalling categories (Fig. 2E) which tend to be maintained in day 35 resistant cells (Fig. 2G) and is emphasised by the expression profiles of genes like *DGAT2*, *EPHX2*, *IDH1* and *PRKAB2* (Supplementary Fig. S2F). The changes in gene expression categories as cells become resistant are further emphasised by comparing GO terms relative to the parental OE19 cells, although this comparison demonstrates that the cell cycle is still suppressed at day 35 whereas various lipid metabolic processes are rapidly altered and maintained at higher levels throughout the time course (Supplementary Fig. S2G). Ingenuity pathway analysis (IPA) revealed the expected inhibition of the upstream regulator ERBB2 but also other cell cycle regulators like MYC and FOXM1 in resistant cells (Supplementary Fig. S2H). Collectively these data reveal that a substantial rewiring of the transcriptome occurs as cells progress to becoming resistant with initial decreases in cell cycle gene expression, accompanied by the sustained activation of metabolic processes.

**Figure 2.**
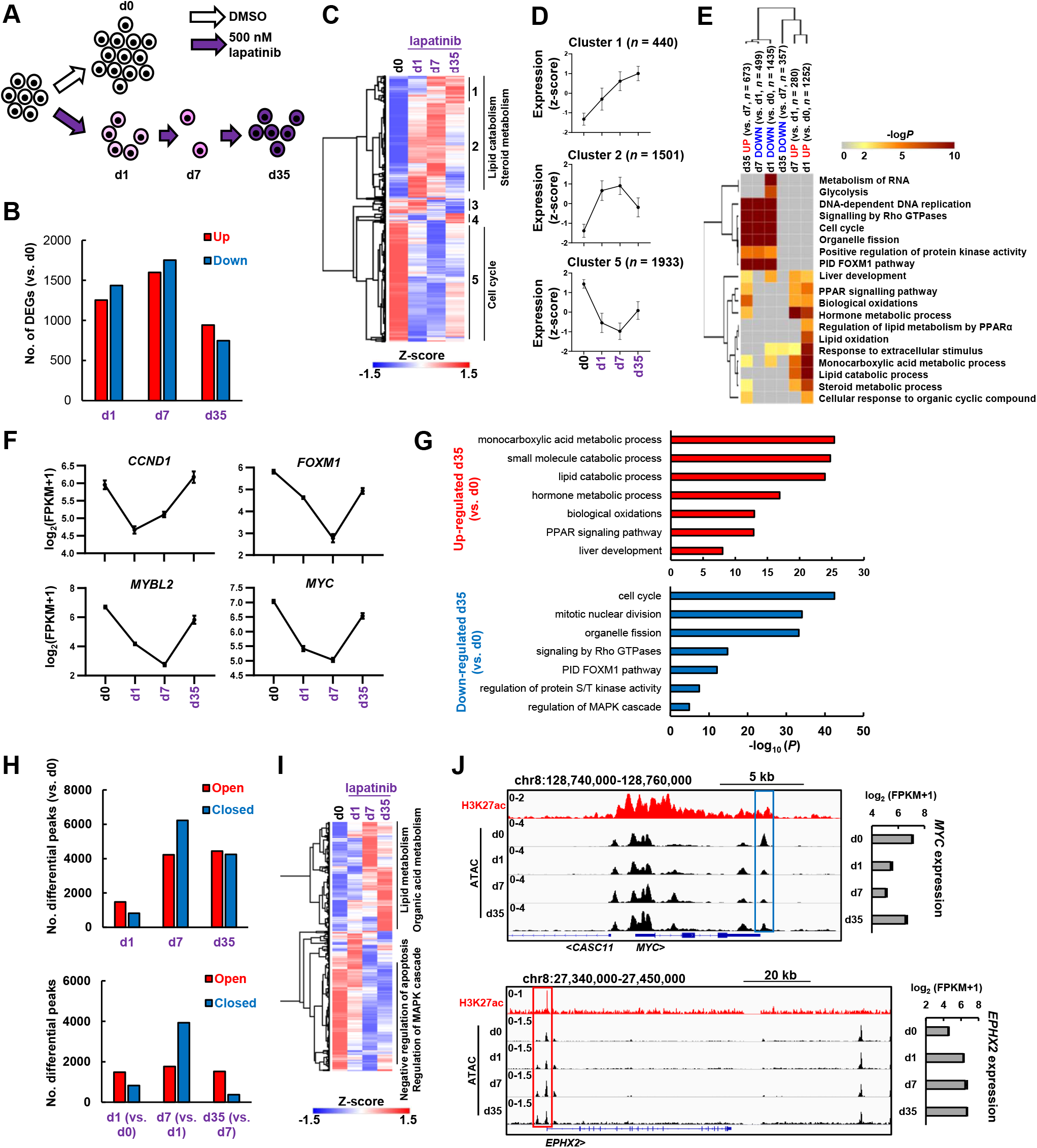
ERBB2 inhibition causes global changes to the transcriptome and chromatin landscape of OAC cells. (A) Schematic of experimental outline. OE19 cells were treated with 500 nM lapatinib for the indicated number of days (d0, d1, d2 and d35). ATAC- and RNA-seq were performed at these timepoints. (B) The number of differentially expressed genes (DEGs) relative to d0 control for the indicated timepoints. DEGs were defined by: 2X linear fold change, FDR < 0.05, FPKM > 1. (C) Heatmap of DEG expression across the timecourse. Clusters of similarly responding genes and associated GO terms are indicated. (D) Average expression profiles of genes in the indicated clusters from (C) across the lapatinib treatment time course. Error bars depict SD. (E) GO analysis using Metascape of DEGs relative to the previous timepoint. (F) Expression of the indicated cell cycle genes across the indicated time points. (G) GO analysis of DEGs in resistant cells (d35) relative to control (d0) cells. (H) The number of all differentially accessible peaks (2X linear fold change, FDR < 0.05) at the indicated timepoints relative to control (d0) cells (top panel) or the previous timepoint (bottom panel). (I) Heatmap showing ATAC-seq signal of differentially accessible chromatin across the timecourse. Prominent GO terms for the major clusters are indicated. (J) Genome browser view of the *MYC* and *EPHX2* loci, highlighting differentially closed and open chromatin respectively after lapatinib treatment. mRNA expression over the timecourse is shown on the right. H3K27ac ChIP-seq (Chen et al., 2020) data is also shown.

To further probe the gene regulatory changes and uncover potential regulatory pathways, we next performed ATAC-seq to assess changes to the accessible chromatin landscape across the same time points towards a lapatinib resistant sate. Widespread changes were observed in the accessible chromatin landscape, which are initiated after 1 day and result in thousands of differentially accessible regions after 7 and 35 days (Fig. 2H; Supplementary Fig. S2I and J; Supplementary Table 2). The majority of the differentially accessible regions are found in intra and intergenic regions that potentially represent potential enhancers (Supplementary Fig. S2K and L). There is a notable increase in promoter accessibility after 7 days whereas there is a tendency for putative enhancer regions to close (Supplementary Fig. S2L). Two broad groups of chromatin regions can be identified, which show general opening or closing across the time course (Fig. 2I) as exemplified by the *MYC* and the *EPHX2* loci which mirrors changes in their gene expression (Fig. 2J). Indeed, more broadly, both promoters and enhancer regions show correlations between changes in accessibility and associated putative target gene expression (Supplementary Fig. S2M). Consistent with transcriptional changes, each set of regions is associated with genes linked to different biological processes, chiefly lipid metabolism and other metabolic processes for regions with increased accessibility and regulation of apoptosis and MAPK cascades for those closing (Fig. 2I; Supplementary S2N). The latter observation is consistent with the expected effects of ERBB2 inhibition.

Collectively, these data demonstrate widespread dynamic changes to the accessible chromatin landscape and associated transcriptome as cells adapt to inhibition of ERBB2 signalling through lapatinib treatment.

### The remodelling of the accessible chromatin state is reversible

Having established the accessible chromatin landscape in lapatinib resistant cells, we next asked whether this is a stable state potentially promoted through the selection of rare mutational events or through the establishment of fixed epigenetic states in the cell population. We therefore profiled the open chromatin landscape of OE19 cells which had been cultured in lapatinib for 35 days (hence adopting a resistant state) followed by drug withdrawal at several timepoints between 1 and 14 days (Fig. 3A). Phosphorylation levels of ERBB2 and downstream ERK and AKT were re-established after 1-2 days of drug withdrawal, demonstrating reinstatement of ERBB2 signalling activity (Fig. 3B). Consistent with this, after 14 days drug withdrawal, the cells again became sensitive to lapatinib re-addition (Fig. 3C).

**Figure 3.**
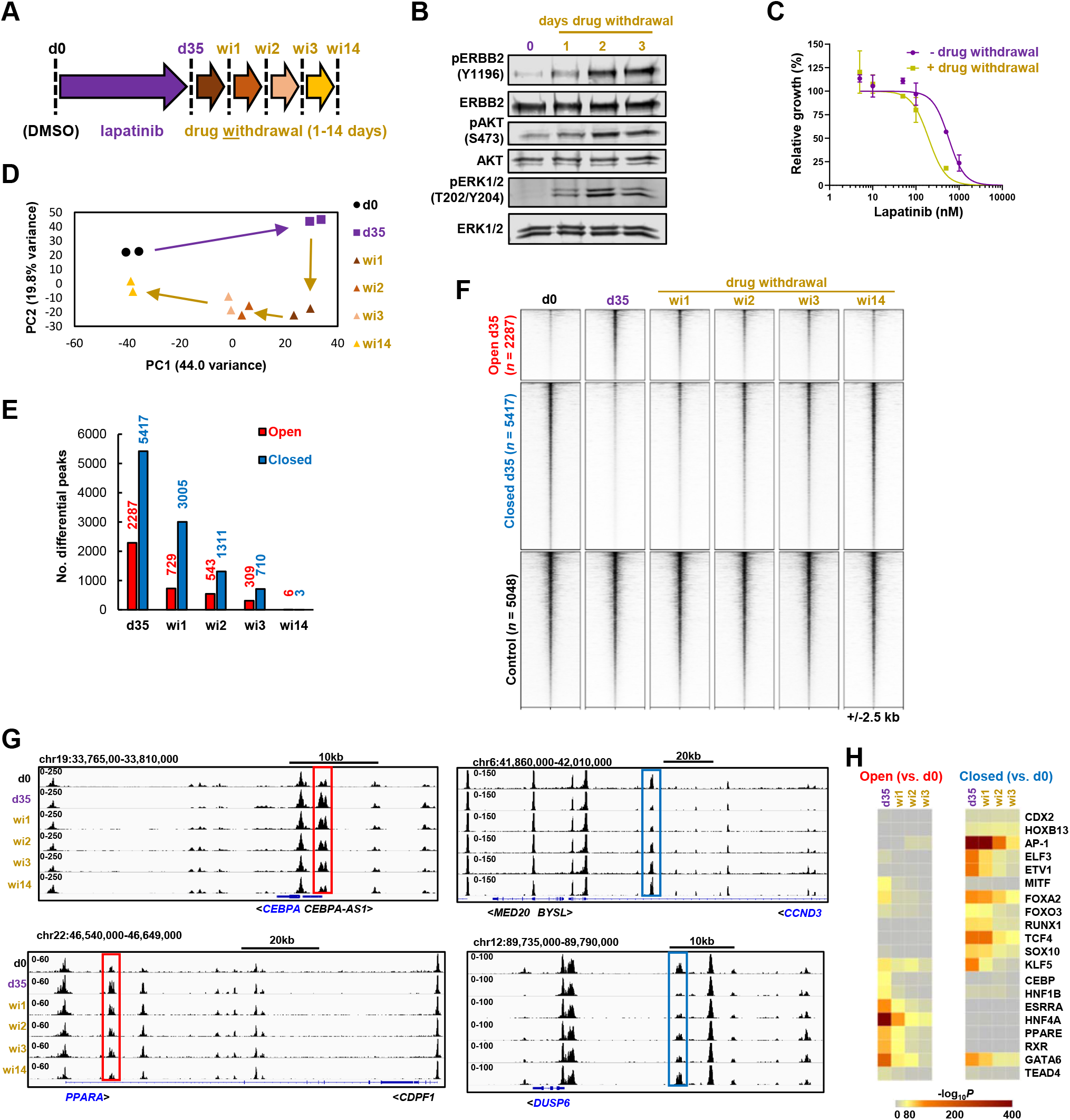
Chromatin accessibility changes in resistant cells are reversible following drug withdrawal. (A) Schematic of experimental outline. OE19 cells were treated with lapatinib for 35 days and then lapatinib was withdrawn. ATAC-seq was performed at the indicated timepoints. (B) Western blot of resistant (d35) OE19 cells following drug withdrawal. (C) MTS growth assay of resistant OE19 cells (d35 lapatinib + d14 lapatinib) (n=2) and drug withdrawal OE19 cells (d35 lapatinib + d14 drug withdrawal)(n=3). Cells were seeded in 500 nM lapatinib and then treated with the indicated doses of lapatinib 24 hours after seeding for 72 hours. Data are shown relative to the lowest dose of drug used. (D) Principal component analysis of OE19 ATAC-seq data of control (d0), resistant (d35 lapatinib) and drug withdrawal cells (wi1,2,3,14). (E) The number of differentially accessible peaks at the indicated timepoints relative to control (d0) OE19 cells. (F) Heatmap of differentially accessible chromatin after d35 lapatinib treatment relative to control cells. (G) Genome browser view of ATAC-seq data in OE19 cells at the indicated timepoints. Differentially accessible peaks that revert following drug withdrawal are highlighted. (H) Heatmap of *P*-values of known motifs enriched in differentially open or closed chromatin at the indicated timepoints (d35 lapatinib treatment and 1-3 days drug withdrawal from this timepoint) relative to control (d0) untreated OE19 cells.

PCA analysis showed a gradual drift of the open chromatin state of resistant cells back towards the parental OE19 cells (Fig. 3D; Supplementary Fig. S3A) which was further supported by Pearson’s correlation analysis which clustered the parental OE19 cells with the resistant cells which have been released from drug treatment for 14 days (Fig. S3B). Furthermore, when we examined the fate of the peaks which changed accessibility in the resistant cells, there was a rapid decay in number after drug withdrawal until barely any differences were detected between the parental cells and the resistant cells after 14 days (Fig. 3E and F). This is exemplified at the locus-specific level where closed chromatin is re-established around the *CEBPA* and *PPARA* loci whereas chromatin reopening is reinstated at the *CCND3* and *DUSP6* loci (Fig. 3G). A number of transcription factor motifs are enriched in peaks showing differentially increased or decreased accessibility after 35 days of drug treatment (see Fig. 4B and C). Interrogation of these enriched transcription factor motifs showed a gradual loss of binding sites for HNF4, GATA and CEBP factors in peaks exhibiting increased accessibility relative to parental cells following drug withdrawal from d35 cells (Fig. 3H, left). Conversely, there was reduced enrichment of the motifs for SOX, AP1 and ETS (ELF3 and ETV1) transcription factors in regions exhibiting decreased accessibility relative to parental cells following drug withdrawal (Fig. 3H, right). Thus, the underlying transcriptional regulatory networks are of OE19 cells are also rapidly re-established following cessation of drug treatment.

**Figure 4.**
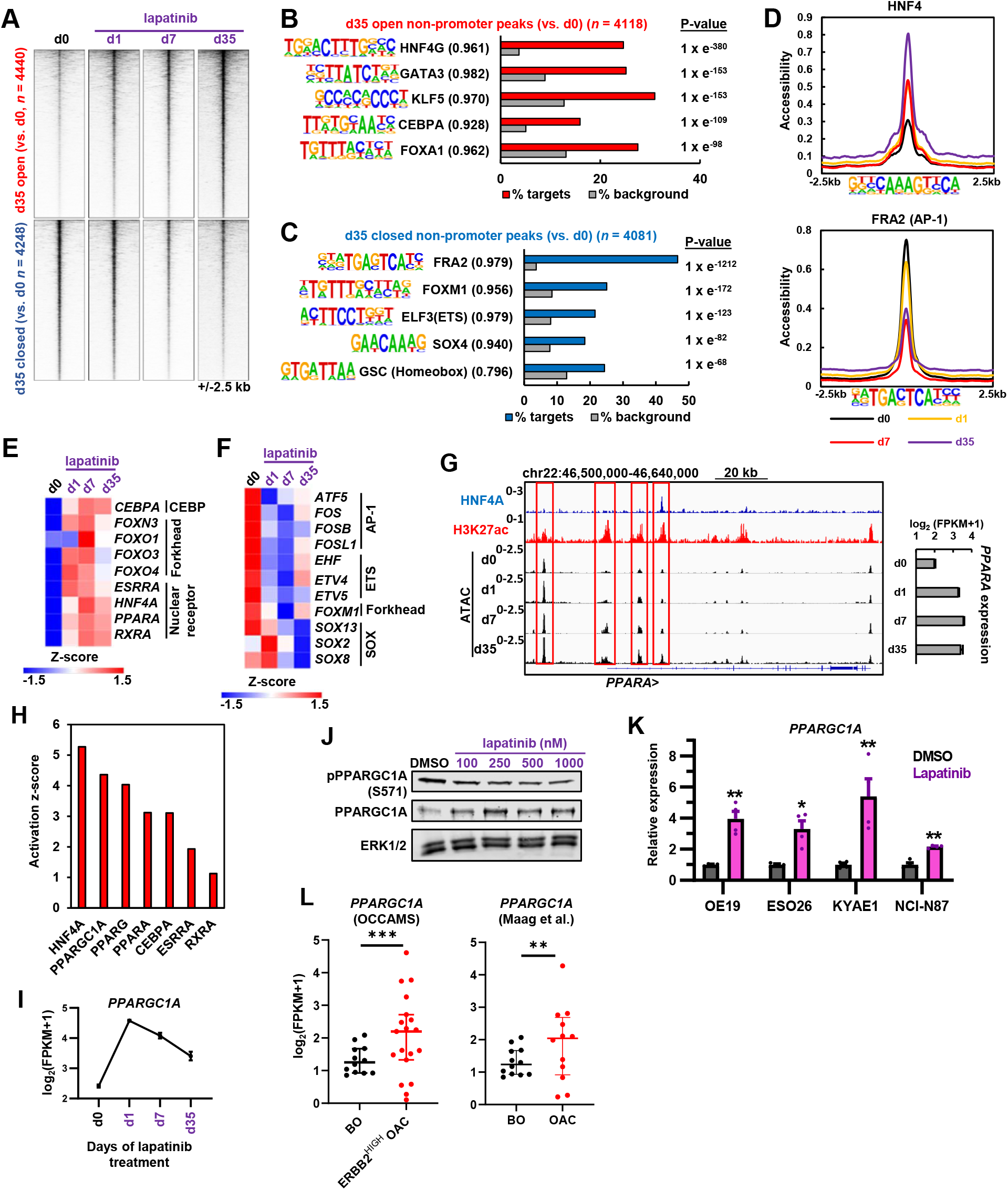
A gastro-intestinal transcription factor programme is activated after ERBB2 inhibition in OAC cells. (A) Heatmap of differentially accessible chromatin regions after 35 days lapatinib treatment (d35) relative to control (d0) cells. Data is also shown for day 1 (d1) and day 7 (d7). (B and C) *De novo* motif enrichment in differentially open (B) or closed (C) non-promoter regions. Motif match score to called transcription factor is shown in brackets. Note that FRA2 is an AP1 family member. (D) Tag density plot of ATAC-seq data across the lapatinib treatment timecourse (d0-d35). Differentially accessible peaks were centred on the HNF4 or AP1 motif. (E and F) Heatmap showing gene expression of up-(E) or down-(F) regulated transcription factors that may bind to motifs enriched in differentially accessible chromatin shown in B and C. Days (d) of lapatinib treatment are shown. Transcription factors are grouped according to their families. (G) Genome browser view of the *PPARA* locus in OE19 cells showing ATAC-seq, HNF4A ChIP-seq (Rogerson et al., 2019) and H3K27ac ChIP-seq (Chen et al., 2020) data. Differentially open peaks are highlighted using red rectangles. Bar chart depicts *PPARA* mRNA expression over the lapatinib treatment timecourse. (H) Ingenuity pathway analysis predicting activated upstream regulators from DEGs in resistant OE19 cells (d35) relative to control (d0) cells. (I) Expression of *PPARGC1A* (PGC1α) over the lapatinib treatment timecourse in OE19 cells. (J) Western blot analysis of PPARGC1A expression and phosphorylation at S571 in OE19 cells treated with the indicated concentrations of lapatinib. (K) RT-qPCR analysis of *PPARGC1A* expression after 24 hours of 500 nM lapatinib treatment. * *P* < 0.05, ** *P* < 0.01, paired t-test, *n* = 3. (L) RNA-seq data showing the expression of *PPARGC1A* in BO (Maag et al., 2017) and OAC (OCCAMS and Maag et al., 2017, as indicated) tissue. ** *P* < 0.01, *** *P* < 0.001.

Taken together these data show a rapid reactivation of the ERBB2 signalling pathway and almost complete reversion of the accessible chromatin landscape to the parental form following drug withdrawal. This argues against selection of a cellular subclone with a unique chromatin landscape and instead demonstrates that the underlying chromatin landscape is reprogrammed based on the intrinsic regulatory pathways available to the OAC cells, and that this is fully reversible upon re-activation of ERBB2 signalling.

### Different transcriptional regulatory pathways are associated with chromatin opening and closing events

Having established the changes in chromatin accessibility during acquisition of resistance to ERBB2 inhibition, we next harnessed this data to uncover the transcription factor networks affected. To focus on the resistance mechanisms, we concentrated on putative enhancer elements (i.e. non-promoter regions) which showed increased or decreased accessibility in cells treated with lapatinib for 35 days. More accessible regions already showed evidence of opening after 1 day of treatment whereas less accessible regions only closed substantially after 7 days (Fig. 4A; Supplementary Fig. S4A). Opening regions showed enrichment of motifs for a range of transcription factors which drive a gastro-intestinal programme in OAC including HNF4, GATA, FOXA, KLF and CEBP factors after 35 days (Fig. 4B; Supplementary Table 2; Rogerson et al., 2019; 2020; Chen et al., 2020; Ma et al., 2021; Pan et al., 2020). These events were already initiated after 1 day treatment (Supplementary Fig. S4F). In contrast, closing regions contained over represented motifs for SOX transcription factors and members of the AP1 and ETS transcription factor families that we previously linked to OAC (Fig. 4C; Supplementary Table 2; Keld et al., 2010; Britton et al., 2017). Closer examination of chromatin accessibility at HNF4A and AP1 motifs further emphasised the dynamic nature of the accessibility around these transcription factor binding sites (Fig. 4D). To examine whether any transcription factors that bind to these motifs exhibit changes in expression, we examined our RNA-seq data and found congruency between directionality of expression and motif presence (Fig. 4E and F). HNF4A, PPARA and several other nuclear hormone receptors show upregulation following lapatinib treatment, alongside CEBPA and several FOX family members (Fig. 4E). Similar increases in expression of the majority of these genes were observed upon depletion of ERBB2 (Supplementary Fig. S4B). Conversely, several SOX and ETS transcription factors (including ETV4/5), and AP1 proteins from the FOS subfamily are downregulated (Fig. 4F). *PPARA* upregulation is associated with the opening of multiple potential regulatory regions throughout its locus (Fig. 4G) whereas *SOX13* downregulation coincides with the closing of several regions (Supplementary Fig. S4C).

We also asked whether these transcription factor networks were similarly affected in other OAC/GAC cell lines containing ERBB2 amplifications and focussed on the initial responses to lapatinib treatment for 1 day because an increase in accessibility centred on the TF motifs like HNF4 was already apparent after 1 day of treatment (Fig. 4D; Supplementary Fig. S4F). There was a moderate overlap in their overall accessibility profiles (Supplementary Fig. S4D) and in the identities of the specific peaks showing differential accessibility following lapatinib treatment (Supplementary Fig. S4E). In all cell lines, opening regions showed enrichment of binding motifs for HNF4A and FOX transcription factors whereas closing regions exhibited enrichment of AP1 motifs (Supplementary Fig. S4F). Thus, the changes to the transcriptional regulatory pathways that we observe in response to drug treatment are generally detected in a range of OAC cell lines.

To further examine which upstream pathways are operational in resistant cells, we performed IPA analysis on genes which were differentially activated after 35 days of lapatinib treatment. Consistent with our transcription factor motif analysis, HNF4A was identified as the most significant upstream regulator alongside a range of other nuclear hormone receptors and the co-activator PPARGC1A/PGC1α (Fig. 4H). Due to its known role as a co-activator for HNF4A (Yoon et al., 2001; Rhee et al., 2003; Charos et al., 2012) and other nuclear hormone receptors like ESRRA and PPARA/G (Mootha et al., 2004; Vega et al., 2000), we further investigated *PPARGC1A* and found that its expression was rapidly induced upon ERRB2 inhibition in OE19 cells and was subsequently maintained well above basal levels in resistant cells (Fig. 4I and J). We also detected a lapatinib-induced decrease in PPARGC1A phosphorylation at S571 (an inhibitory phosphorylation event; Li et al., 2007), providing further activation potential. Similarly, lapatinib treatment caused increased *PPARGC1A* expression in a range of other OAC/GAC lines (Fig. 4K). Moreover, *PPARGC1A* levels are already elevated in samples from patient OAC tumours containing high *ERBB2* levels compared to the precursor Barrett’s oesophagus (Fig. 4L). Several nuclear hormone receptors showed changed expression in tumours with high levels of *ERBB2* with some being elevated (*ESRRA*) whereas others were either unchanged (*HNF4A*) or showed reduced levels (*PPARA*) (Supplementary Fig. S4G). There is therefore no uniform response to ERBB2 amplification in patients. Thus, a simple model whereby ERBB2 signalling directly suppresses nuclear hormone receptor expression and activity relative to the precursor Barrett’s state does not appear to be operational.

Collectively these data demonstrate that the lapatinib resistant state is characterised by the activation of a transcription factor network that comprises a set of gastro-intestinal transcription factors including HNF4A, and additional nuclear hormone receptors alongside the coactivator protein PPARGC1A.

### HNF4A activity is enhanced and required for lapatinib resistance

To provide further mechanistic insights into how transcriptional regulatory networks influence the lapatinib resistant cell state we first focussed on the nuclear hormone receptor HNF4A due to the enrichment of its binding motifs in accessible regions appearing in resistant cells (Fig. 4B). We performed ChIP-seq analysis to ask whether HNF4A gained additional activities through changing its targets following lapatinib treatment (Supplementary Fig. S5A and B; Supplementary Table 3). We first examined the HNF4A signal at chromatin loci that increased in accessibility one day after lapatinib treatment and found a general increase in HNF4A binding occurred after lapatinib treatment (Fig. 5A, top; Supplementary Fig. S5C). This is consistent with the increases in HNF4A expression we observe (Fig. 4E). Furthermore, this increased binding was accompanied by increased levels of the activate histone mark H3K27ac, consistent with an activating event. In contrast, closing regions showed only low levels of HNF4A binding in parental cells with little change in binding triggered by lapatinib treatment, although inactivation was suggested by the decreases in H3K27ac (Supplementary Fig. 5A, bottom; Supplementary Fig. S5C). We also partitioned the HNF4A binding regions into those with increased (ERBB2 suppressed) or decreased (ERBB2 activated) occupancy in lapatinib treated cells (Supplementary Table 3). HNF4A peaks which show increased occupancy also exhibited increases in both H3K27ac levels and chromatin opening following lapatinib treatment (Fig. 5B; top) whereas HNF4A peaks showing reduced occupancy also demonstrated reduced levels of acetylation but little change in chromatin opening (Fig. 5B; bottom). These results indicate that elevated levels of HNF4A binding in response to lapatinib treatment are associated with chromatin activation events and these accompany gene expression changes as exemplified by the *IMPA1* (Fig. 5C), *KDM5B* (Supplementary Fig. S5D), loci. Indeed, there is a more widespread strong correlation between the changes in HNF4A binding activity and the changes of H3K27ac observed following lapatinib treatment (Supplementary Fig. S5E). Further analysis of genes neighbouring HNF4A binding regions that exhibit increased occupancy in response to lapatinib treatment, showed that these activation events correlate with increased gene expression under the same conditions (Fig. 5D). These HNF4A binding regions are rich in HNF4A binding motifs (70% of regions), consistent with direct binding but we also observed enrichment for GATA motifs suggesting combinatorial control at a subset of sites (Fig. 5E). In contrast, in regions showing decreased HNF4A occupancy, HNF4A binding is less associated with HNF4A motifs (25% of regions) and instead, AP1 is the highest ranked motif, suggesting potential indirect recruitment of HNF4A to these regions (Fig. 5F). Functionally, regions which both opened (d1), and harboured HNF4A binding regions (d2) following lapatinib treatment are close to genes associated with GO terms encompassing a variety of mitochondrial related and metabolic processes, consistent with a potential role in metabolic reprogramming (Supplementary Fig. S5F). To establish the relevance of these regulatory events in drug resistance, we depleted HNF4A (Supplementary Fig. S5G) and monitored the re-emergence of cell proliferation following lapatinib treatment. Loss of HNF4A blunted the re-establishment of cell growth and resistance in OE19 (Fig. 5G) cells. In contrast, HNF4A loss did not cause reduced growth of parental OE19 cells in the absence of lapatinib treatment (Supplementary Fig. S5H).

**Figure 5.**
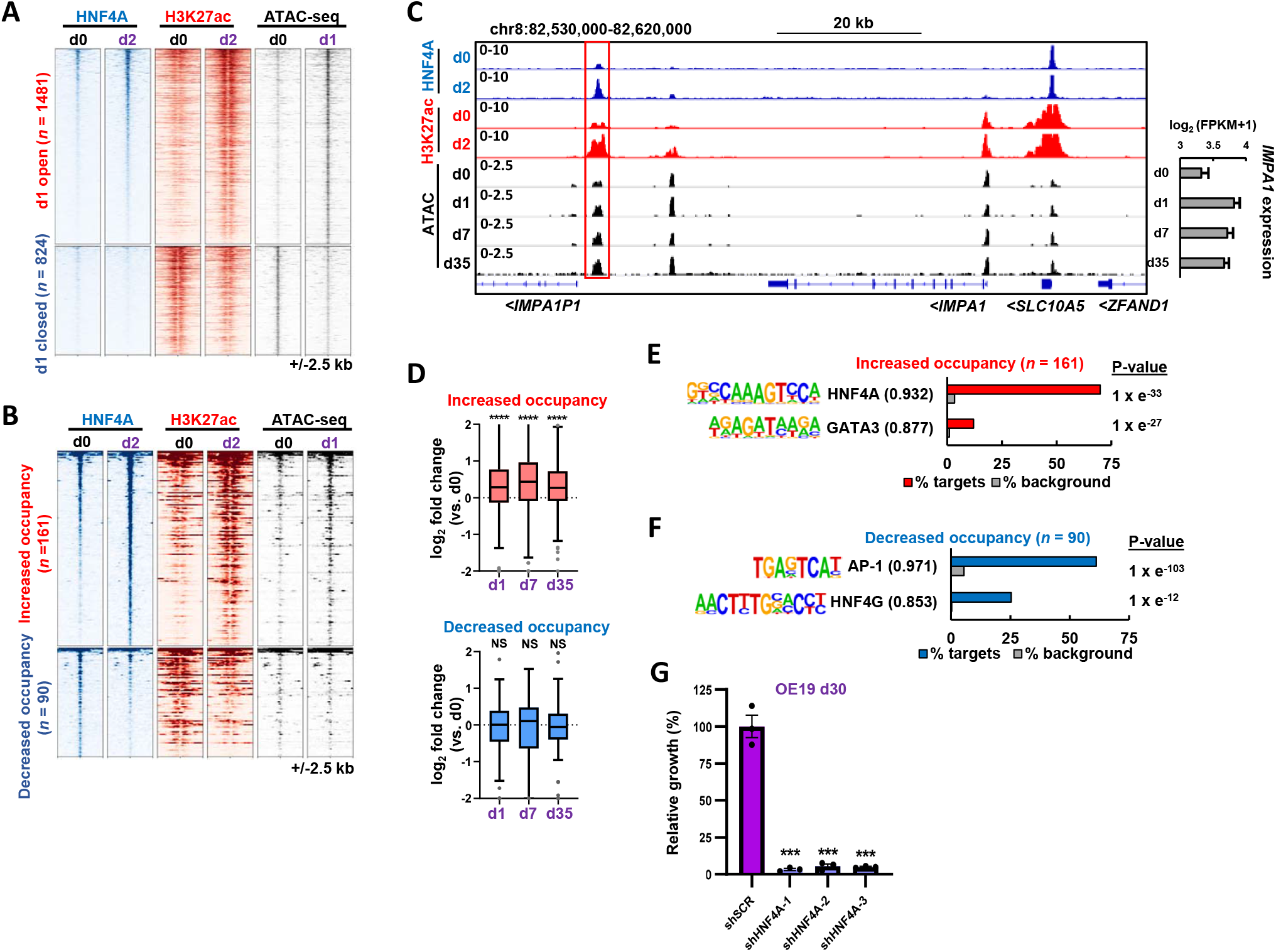
HNF4A regulatory activity is enhanced following ERBB2 inhibition. (A) Heatmap of differentially open or closed ATAC-seq peaks in OE19 cells after 1 day 500 nM lapatinib treatment (d1) relative to control cells (d0)(right). HNF4A and H3K27ac ChIP-seq data are shown in control (d0) cells or after 2 days 500 nM lapatinib treatment (d2)(left and centre). (B) Heatmap showing HNF4A binding sites with increased (ERBB2 suppressed) or decreased (ERBB2 enhanced) HNF4A binding after 2 days lapatinib treatment (d2) relative to control cells (d0). (C) Genome browser view of the *VEGFA* locus in OE19 cells. Red box highlights a putative enhancer in which HNF4A binding increases after 2 days lapatinib treatment. *VEGFA* mRNA expression over the lapatinib time course is shown on the right. (D) Fold change in expression of genes relative to control cells (d0) of genes annotated to differentially bound HNF4A sites using the basal plus extension model (GREAT). **** *P* < 0.0001, NS – not significant; Wilcoxon matched pairs signed rank test for each timepoint relative to d0. (E and F) *De novo* motif enrichment in differentially bound HNF4A binding sites after 2 days lapatinib treatment. HNF4A binding sites exhibiting enhanced (E) or decreased (F) occupancy following lapatinib treatment are shown. Motif match score to called transcription factor is shown in brackets. Sites included both promoter and non-promoter peaks. (G) Crystal violet growth assay of OE19 cells treated with control shRNA (shSCR) or shRNA targeting HNF4A (shHNF4A). Cells were treated with 500 nM for 30 days and data are shown relative to shSCR control, *** *P* < 0.001, unpaired T-test, *n* = 3, error bars depict SEM.

These results therefore demonstrate that ERBB2 inhibition triggers enhanced HNF4A binding to chromatin which is associated with chromatin and gene activation events. These events are important for the emergence of resistance, potentially through metabolic reprogramming.

### PPARGC1A promotes metabolic changes during acquisition of the resistant state

To further understand the transcription factor networks driving the resistant state, we turned to PPARGC1A, which has previously been shown to act as a potential coactivator protein for HNF4A (Yoon et al., 2001; Rhee et al., 2003; Charos et al., 2012). Our results have implicated PPARGC1A in driving the chromatin changes in the resistant state (Fig. 4H), therefore we performed ChIP-seq in OE19 cells to identify the binding sites for PPARGC1A (Supplementary Fig. S6A; Supplementary Table 4). Unexpectedly, we found that PPARGC1A and HNF4A exhibited virtually mutually exclusive binding locations (Fig. 6A; Supplementary Fig. S6B and C). These two proteins therefore likely contribute to lapatinib resistance through different regulatory activities and target genes.

**Figure 6.**
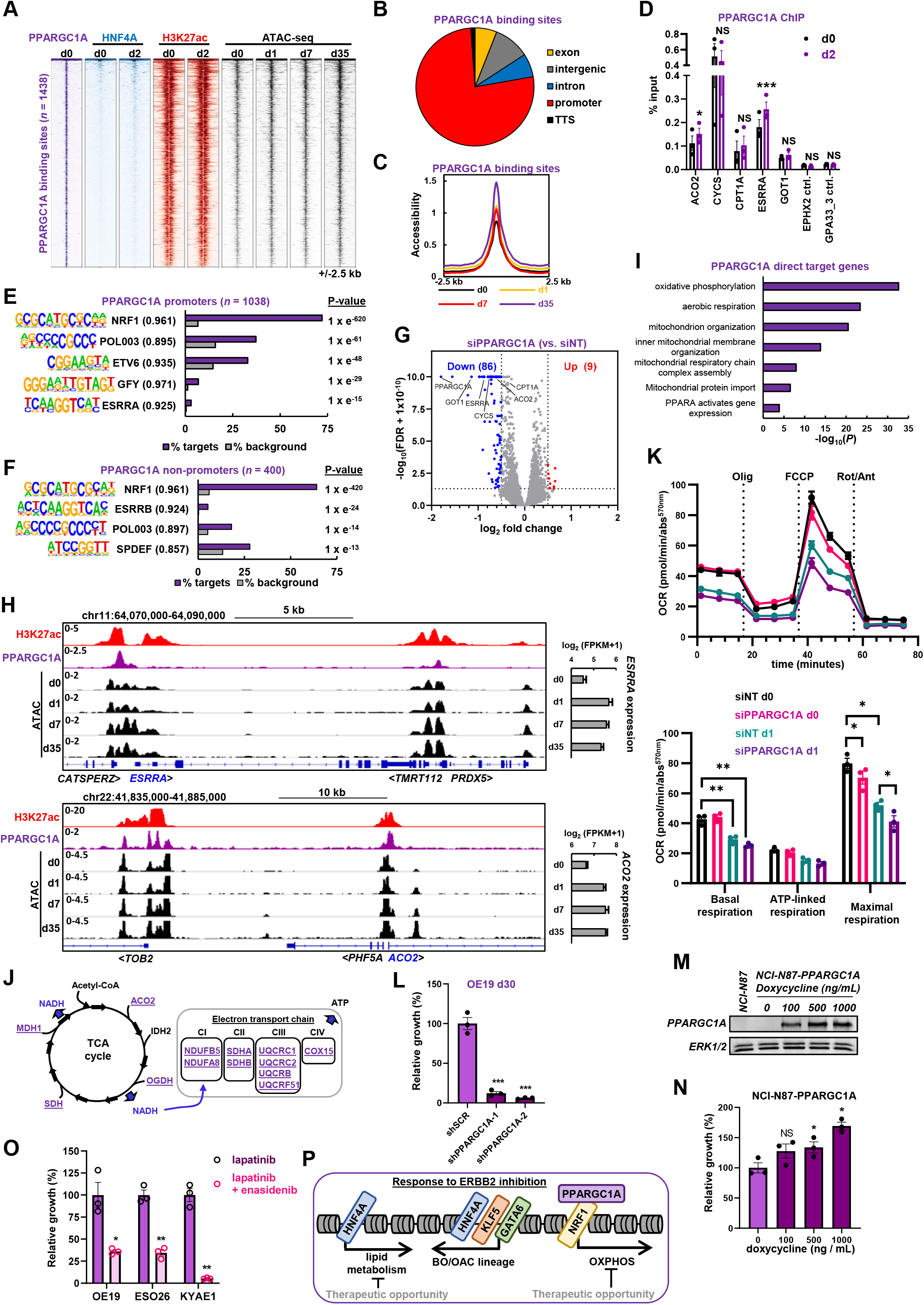
PPARGC1A regulates mitochondrial processes and supports the emergence of resistance. (A) Heatmap of PPARGC1A binding sites in OE19 cells and HNF4A binding, H3K27ac and ATAC-seq signal in control (d0) cells or after lapatinib treatment for the indicated number of days (d1-d35). (B) Genomic distribution of PPARGC1A binding sites in OE19 cells. (C) Tag density plot of chromatin accessibility over the lapatinib timecourse at PPARGC1A binding sites. (D) PPARGC1A ChIP-qPCR in OE19 cells treated with DMSO (d0) or with 500 nM lapatinib (d2) for 2 days (d0 or d2). Statistical significance was determined using a paired T-test; *\*\*\* P* < 0.001, * *P* < 0.05, NS – non-significant, *n* = 3. (E and F) *De novo* motif enrichment at PPARGC1A-bound promoters (E) or non-promoter regions (F). Motif match score to called transcription factor is shown in brackets. (G) Volcano plot highlighting DEGs after OE19 cells were transfected with siPPARGC1A and treated with 500 nM lapatinib for 24 hours. Significantly down- or up-regulated genes are indicated in blue and red respectively and were defined by log2fold change > 0.5 and FDR < 0.05. (H) Genome browser view of the *ESRRA* and *ACO2* loci and surrounding genomic region in OE19 cells. ATAC-seq, PPARGC1A binding and H3K27ac signal is shown. Gene expression from RNA-seq data is shown on the right. Duration of lapatinib treatment in days (d) is shown. (I) GO analysis using Metascape of PPARGC1A direct target genes (*n* = 123) in OE19 cells treated with lapatinib for 24 hours. Direct target genes were defined as genes down-regulated (FDR < 0.05) by siPPARGC1A that were also annotated to a PPARGC1A ChIP-seq peak using the basal plus extension model (GREAT). (J) PPARGC1A activated direct target genes (blue, underlined) are shown on the TCA cycle and the associated electron transport chain. (K) Mitochondrial stress test in OE19 cells treated with siPPARGC1A and DMSO or 500 nM lapatinib for 24 hours. OCR – oxygen consumption rate. Statistical significance was determined using a paired t-test; * *P* < 0.05, ** *P* < 0.01, *n* = 3. (L) Crystal violet growth assay of OE19 cells treated with control shRNA (shSCR) or shRNA targeting PPARGC1A (shPPARGC1A). Cells were treated with 500 nM lapatinib for 30 days and data are shown relative to shSCR control. *** *P* < 0.001, unpaired T-test, *n* = 3. (M) Western blot of PPARGC1A levels in parental NCI-N87 or stable NCI-N87 expressing inducible PPARGC1A (NCI-N87-PPARGC1A). Cells were treated with the indicated amounts of doxycycline for 48 hours. (N) Crystal violet growth assay of NCI-N87-PPARGC1A cells treated with 250 nM lapatinib and the indicated doses of doxycycline for 72 hours. (O) Crystal violet growth assay of the indicated cell lines treated with lapatinib or lapatinib plus 5 μM enasidenib. OE19 cells were treated with 500 nM lapatinib for 35 days, ESO26 cells (*PIK3CA* activating mutation) were treated with 100 nM lapatinib and 250 nM MK-2206 (AKTi) for 18 days and KYAE1 cells were treated with 100 nM lapatinib for 28 days. * P < 0.05, ** P < 0.01, paired T-test, n = 3. (P) Model showing the changes in transcription factor activity elicited by ERBB2 inhibition on top of the programme driving the BO/OAC phenotype.

Next, we analysed PPARGC1A in more detail to uncover its regulatory activities. PPARGC1A binding was detected at 1,438 sites that are largely associated with promoter regions (Fig. 6B). These regions exhibited a general increase in accessibility upon lapatinib treatment, which was most apparent at day 35 (Fig. 6A and C; Supplementary Fig. S6D). In addition, the PPARGC1A binding regions are associated with the active histone mark H3K27ac, in both parental and lapatinib treated cells (Fig. 6A) and exhibit slight increases in acetylation after drug treatment (Supplementary Fig. S6D). We tested a panel of these binding sites in lapatinib treated cells and observed only moderate increases in PPARGC1A occupancy, although significant increases were found at the *ACO2* and *ESRRA* loci (Fig. 6D). However, reductions in inhibitory phosphorylation events on PPARGC1A were observed indicating a likely increase in pre-bound PPARGC1A activity (Fig. 4J). To gain insights into the potential transcription factors that might recruit PPARGC1A, we searched for enriched DNA motifs and uncovered the NRF1 motif in both promoters and potential distal regulatory regions in the vast majority of binding regions (Fig. 6E and F). In keeping with the lack of overlap with the HNF4A ChIP-seq data, we did not observe co-occurrence of the HNF4A binding motif. We took advantage of ChIP-seq data for NRF1 from other cell lines and observed a large overlap with PPARGC1A binding regions (Supplementary Fig. S6B). NRF1 has previously been implicated in controlling mitochondrial function and its metabolic activities (Evans et al., 1990; Wu et al., 1999; Cam et al., 2004) and consistent with this, two of the top GO terms for genes associated with PPARGC1A binding regions are “mitochondrial organisation” and “TCA cycle and respiratory electron transport” (Supplementary Fig. S6E).

To uncover the regulatory consequences of PPARGC1A binding, we next depleted PPARGC1A (Supplementary Fig. S6F and G) and performed RNA-seq on OE19 cells after lapatinib treatment for 24 hrs (Supplementary Fig. S6H). PPARGC1A depletion mainly led to downregulation of gene expression with 86 genes exhibiting significantly reduced levels (Fig. 6G; 1.4 fold change, FDR < 0.05; Supplementary Table 4). By analysing all significantly downregulated genes, we observed a large number of direct targets for PPARGC1A that are associated with PPARGC1A binding peaks (Supplementary Fig. S6I; left; Supplementary Table 4) including *ESSRA* and *ACO2* (Fig. 6H). The number of direct target genes increased following lapatinib treatment (Supplementary Fig. S6I; right; Supplementary Table 4). Many of these directly activated targets (36.6%) show upregulation in response to lapatinib treatment as exemplified by *ESSRA* and *ACO2* (Fig. 6H; Supplementary Fig. S6J). GO term analysis of the directly activated PPARGC1A target genes uncovered several terms associated with mitochondrial function and aerobic respiration (Fig. 6I) and many of these genes are key components of the TCA cycle or the electron transport chain (Fig. 6J). We therefore examined mitochondrial activity by measuring the oxygen consumption rate in parental OE19 cells and cells treated with lapatinib in the presence and absence of PPARGC1A depletion (Fig. 6K). Basal and maximal respiration rates were dampened by lapatinib treatment and maximal respiration was further reduced by PPARGC1A depletion. In contrast, the glycolytic rate as assessed by the extracellular acidification rate, was unaffected by PPARGC1A depletion (Supplementary Fig. S6K). However, a lapatinib-mediated decrease in glycolytic rate was observed, suggesting a general decrease in metabolic activity.

Having established that the PPARGC1A-driven pathway is one of the predominant transcriptional pathways activated in OAC cells following lapatinib treatment, we asked whether this transcriptional regulator plays a role in acquired resistance. We created two different shRNA constructs to deplete PPARGC1A (Supplementary Figs. S5G and S6L) and tested the re-emergence of cell growth following extended treatment with lapatinib. Control OE19 cells transduced with scrambled shRNA constructs restarted proliferation after 30 days lapatinib treatment, but cells depleted of PPARGC1A failed to resume growth (Fig. 6L). In contrast, OE19 cells grown in the absence of lapatinib showed no reductions in growth following PPARGC1A depletion (Supplementary Fig. S5H). Reciprocally, we performed a gain of function experiment in the gastric adenocarcinoma cell line NCI-N87 which harbours an *ERBB2* amplification. This cell line is sensitive to ERBB2 inhibition (Supplementary Fig. S1E) and has a similar open chromatin profile to several OAC cell lines and patient samples (Fig. 1C) but has relatively low levels of *PPARGC1A* expression (Fig. S6M). We inserted a doxycycline inducible *PPARGC1A* transgene (Fig. 6M) and tested growth in the presence of lapatinib. High levels of PPARGC1A promoted enhanced cell growth in the presence of lapatinib (Fig. 6N). To see if we could prevent the emergence of resistance by targeting the metabolic pathways regulated by PPARGC1A, we treated OAC cells with lapatinib plus the IDH2 inhibitor enasidenib. IDH2 is an important enzymatic component of the citric acid cycle and enasidenib has been previously suggested as a mechanism to target PPARGC1A-driven mitochondrial respiration (De Vitto et al., 2019), and whilst enasidenib is more specific to mutant IDH2, it can still inhibit WT *IDH2* (Yen et al., 2017) that is present in parental OE19 cells (Tate et al., 2019). Treatment of OAC cells with enasidenib in the absence of lapatinib only had a slight effect on cell growth (Supplementary Fig. S6N). In contrast, co-treatment of OE19, ESO26 and KYAE1 cells with lapatinib and enasidenib impaired the emergence of resistance (Fig. 6O). Thus, re-purposing of clinically used mitochondrial inhibitors could potentially prevent the emergence of resistance to RTK inhibitors in OAC.

Collectively, our data demonstrate that a PPARGC1A-driven pathway becomes activated by ERBB2 inhibition with lapatinib. Regions bound by this coactivator protein become more accessible and associated target genes are activated following lapatinib treatment, leading to changes in mitochondrial activity and oxidative phosphorylation. This regulatory activity is essential for the resumption of proliferative activity during the acquisition of resistance.

## Discussion

Drug resistance is a major clinical problem for cancer treatment, especially when targeting signalling pathways (reviewed in Boumahdi et al., 2020). This applies to OAC where compensatory mutational changes have been shown to be acquired to permit resistance to the ERBB2 inhibitor trastuzumab (Kim et al., 2014; Janjigian et al., 2018; Wang et al., 2019). However, there are many ways for tumours to acquire resistance, and recently it has been shown that changes to the regulatory chromatin landscape can permit resistance in several scenarios, including endocrine resistance in breast cancer (Bi et al., 2020), BET inhibitor resistance in acute myeloid leukaemia (Bell et al., 2019), and PDGFRA inhibitor resistance in glioblastoma (Liau et al., 2017). Here we have focussed on RTK pathways driven by ERBB2 amplifications in OAC and demonstrate that widespread rewiring of the regulatory chromatin landscape is a major driver towards acquiring drug resistance.

The integration of chromatin changes with transcriptomic changes enabled us to uncover transcriptional regulatory networks that are rewired as cells respond to drug treatment and develop resistance. Shortly after treatment, chromatin closing is instigated which corresponds to transcriptional programmes driven by AP1 and ETS family transcription factors. These families of transcription factors have previously been shown to play important roles in OAC (Keld et al., 2010; Britton et al., 2017; Chen et al., 2020), and lapatinib acts to shut these down. Conversely, chromatin opening causes these programmes to be replaced with a network driven by a set of transcription factors that are usually associated with early intestinal development, typified by HNF4, KLF5, FOXA and GATA factors (Fig. 6P). This set of transcription factors has previously been shown to be operational in both Barrett’s and OAC (Rogerson et al., 2019, 2020; Chen et al., 2020; Ma et al., 2021; Pan et al., 2020) and suggests that ERBB2 activation leads to at least partial suppression of their activity in OAC cells. Alternatively, additional compensatory pathways may be activated in response to drug treatment which further enhance the activity of this set of transcription factors. Further analysis also implicated the transcriptional co-activator PPARGC1A in regulating the gene expression programmes that arise as cells acquire resistance (Fig. 6P). These transcription factors are associated with changes in gene expression changes linked to various metabolic programmes, suggesting that metabolic reprogramming allows cells to persist in the presence of ongoing lapatinib treatment. Our findings in OAC are consistent with work in ERBB2-amplified breast cancer cells where metabolic adaptations to lapatinib treatment occur which shift the cells towards mitochondrial energy metabolism (Deblois et al., 2016). However, in breast cancer these changes are driven by the transcription factor, ERRα demonstrating that resistance to the same inhibitor can occur in two different cancers through reprogamming of different pre-existing gene regulatory networks.

By focussing on HNF4A and PPARGC1A, we provide more detailed analysis of their downstream regulatory activities. PPARGC1A has previously been shown to be a co-activator for several nuclear hormone receptors, including HNF4A (Yoon et al., 2001; Rhee et al., 2003; Vega et al., 2000). However, unexpectedly we found little evidence for chromatin co-occupancy. This is reflected in the metabolic programmes they are implicated in, with PPARGC1A primarily associated with oxidative phosphorylation whereas HNF4A controls various lipid metabolic processes. Interestingly, both converge on programmes controlling mitochondrial functions, suggesting that mitochondrial changes may be key in promoting resistance. Both HNF4A and PPARGC1A activity is required for acquiring resistance, underlying the importance of their downstream regulatory programmes. Indeed, when we inhibit oxidative phosphorylation by targeting IDH2, resistance is impaired, providing more direct evidence that this metabolic switch is important for cancer development. These findings are consistent with studies in other cancer types where PPARGC1A-driven programmes play an important role in drug resistance in melanomas (Haq et al., 2013) and glioblastoma (Zhang et al., 2020). In both cases, PPARGC1A-dependent metabolic reprogramming is elicited following inhibition of either a RTK (MET) or a downstream pathway component (BRaf), demonstrating a common response to RTK pathway inhibition in rewiring metabolic programmes in OAC and these cancers. Furthermore, HNF4A loss in pancreatic cancer also drives metabolic reprogramming and correlates with a metabolic switch to dependence on glycolytic activity (Brunton et al., 2020), suggesting a more widespread role for HNF4A in this context.

After 5 weeks of lapatinib treatment, OAC cells begin to proliferate again despite ERBB2 being inactive, suggesting that other compensatory signalling pathways must be activated which is reflected by the re-activation of ERK. However, this is not a stable state and is fully reversible after drug withdrawal and the open chromatin landscape reverts to its initial state within days. This has important implications for considering treatment regimes, as drug holidays could allow cells to be re-sensitised to ERBB2 inhibition, thereby reducing the selective pressure to select for additional mutational events. Our work has additional therapeutic implications as selective targeting of the persistor cells may prevent transition to a resistant state. One route would be through direct inhibition of HNF4A or PPARGC1A, or alternatively through targeting their regulatory programmes as exemplified by IDH2 inhibition with enasidenib, a clinically approved drug (Yen et al., 2017; Stein et al., 2017).

In summary, we have demonstrated that OAC cells undergo dramatic remodelling of their accessible chromatin landscape following ERBB2 inhibition. This chromatin remodelling revealed that the HNF4A and PPARGC1A regulated transcriptional regulatory networks play a pivotal role in promoting the emergence of drug resistant cancer cells and represent potential therapeutic targets to enhance the efficacy of ERBB2 inhibition strategies. More generally, such an approach could be combined with inhibitors of other RTK pathway components given the high frequency of mutational events in this pathway in OAC (Cancer Genome Atlas Research Network et al, 2017; Frankell et al., 2019).

## Methods

### Cell culture and treatments

OE19, ESO26 and NCI-N87 cells were cultured in RPMI 1640 (ThermoFisher Scientific, 52400) supplemented with 10% foetal bovine serum (ThermoFisher Scientific, 10270) and 1% penicillin/streptomycin (ThermoFisher Scientific, 15140122). KYAE1 cells were cultured in 1:1 RPMI 1640:F12 (Thermo Fisher, 11765054) supplemented with 10% foetal bovine serum and 1% penicillin/streptomycin. HEK293T and HET1A cells were cultured in DMEM (ThermoFisher Scientific, 22320-022) supplemented with 10% foetal bovine serum. CP-A cells were cultured in keratinocyte serum free media (ThermoFisher Scientific, 17005042) supplemented with 10% foetal bovine serum, 5 ng/mL EGF (ThermoFisher Scientific, 17005042), 50 μg/mL bovine pituitary extract (ThermoFisher Scientific, 17005042) and 1% penicillin/streptomycin. Cell lines were cultured at 37°C, 5% CO_2_ in a humidified incubator. For inhibitor treatment, cells were seeded at a density of 2×10^4^ cells/cm^2^. 24 hours after seeding, cells were treated with inhibitors [ERBB2/EGFR, Lapatinib (Selleckchem, S1028); AKT, MK-2206 (Selleckchem, S1078); IDH2, Enasidenib (Selleckchem, S8205) were reconstituted in dimethyl sulfoxide (DMSO)] or vehicle control. For long term treatments, inhibitor and media was replenished every 72 hours. To ensure all timepoints were treated equally, 24 hours prior to the endpoint fresh media/inhibitor was added to cells. For drug withdrawal experiments lapatinib resistant cells were first generated by treatment with lapatinib for 35 days. After 35 days of drug treatment cells were trypsinised, counted and seeded at a density of 2×10^4^ cells / cm^2^ in the continued presence of lapatinib. 24 hours after seeding, media containing lapatinib was removed, cells were washed 3X with PBS to wash out inhibitor before adding fresh media to the cells.

### Organoid culture

WTSI_OESO-009 (CAM408 – Li et al., 2018) was a gift from Rebecca Fitzgerald. Organoids were cultured based on protocols described in Li et al., 2018, except organoids were cultured in IntestiCult™ Organoid Growth Medium (STEMCELL Technologies, 06010). Organoids were cultured in 6-well plates. To passage organoids, media was removed and organoids were washed with PBS. 1 mL PBS was added and organoids were dissociated from BME-2 (Cultrex, 3533-005-02) by pipetting. The organoid suspension was centrifuged at 1000 RCF for 5 minutes and the supernatant was discarded. A single cell suspension was then form by the addition of 1 mL TrypLE™ Express (Gibco, 12609-013), and cells were transferred to a 1.5 mL tube and incubated for 15 minutes, 1000 rpm, 37 °C. Single cells were centrifuged at 1800 RCF rpm for 5 minutes and the supernatant was removed by pipetting. Single cells were re-suspended in 200 μL 10 mg / mL BME-2 and 20 μL cells / BME-2 suspension were then pipetted into a 6-well plate, before incubating for 15 minutes at 37 °C, 5% CO2 to enable BME-2 to polymerise. 1.5 mL IntestiCult™ Organoid Growth Medium was then added to culture the cells.

For MTS growth assays, organoids were formed in 3D culture as described and then dissociated from BME-2 by pipetting. Organoids were then counted using a haemocytometer and 5×10^3^ organoids were seeded into 96-well plates. For ATAC-seq, organoids were dissociated from BME-2 into single cells using TrypLE. ATAC-seq libraries were then generated.

### Cell growth, cell cycle and apoptosis assays

MTS growth assays (Promega, G3580) were performed in 96-well plate format according to the manufacturer’s protocol. Absorbance readings were taken at 490 nm on a SPECTROstar Nano Micoplate Reader (BMG LABTECH).

Crystal violet assays were performed by fixing cells with 4% paraformaldehyde for 10 minutes. Cells were stained using 0.1% crystal violet (Sigma-Aldrich, HT90132) for 30 minutes at room temperature. Plates were rinsed with water and left to dry before solubilising dye in 10% acetic acid for 10 minutes at room temperature with gentle shaking. Absorbance readings were taken at 570 nm on a SPECTROstar Nano Micoplate Reader (BMG LABTECH). Data was uniformly transformed so that the mean of the control sample was represented as 100%.

For cell cycle analysis, media was collected from cells and cells were dissociated using trypsin, using the collected media to quench trypsin. Cells were collected by centrifugation, washed with PBS and then fixed by the addition of 70% ethanol, pre-cooled to −20 °C. Fixed cells were then stored at −20 °C until required. Cells were then pelleted and washed with PBS before the addition of 400 μL 50 μg/mL propidium iodide (Sigma, P4864). Cells were then analysed by the University of Manchester Flow Cytometry Core Facility on a BD Biosciences LSRFortessa™. Data was analysed using ModFit LTTM software to determine the percentage of cells in G0/G1, S or G2/M phase.

For apoptosis assays, 2×10^5^ cells were seeded into 6-well plates. 24 hours later, fresh culture media was added, and cells were treated with 500 nM lapatinib or vehicle control. 30 μM propidium iodide (Sigma, P4864) was added to culture medium to measure apoptosis. Cells were imaged every 20 minutes for 72 hours using an Incucyte ZOOM (ESSEN Bioscience), maintained at 37 °C, 5% CO_2_. The number of apoptotic cells was determined by the number of red fluorescent cells and the data was exported to Prism 8 (GraphPad).

### siRNA transfection

4×10^5^ cells were reverse transfected with 25 pmol siRNA using Lipofectamine™ RNAiMAX transfection reagent (ThermoFisher Scientific, 13778150) according to the manufacturer’s instructions. Cells were seeded into 6-well plates. SMART-pool siRNAs for control non-targeting siRNA (Dharmacon, D-001810-10-0020), siERBB2 (Dharmacon, L-003126-00-0005) and siPPARGC1A (Dharmacon, L-005111-00-0005) were used.

### Lentiviral vectors, production and transduction

The PPARGC1A (PGC1α) gene was amplified from the vector pcDNA myc PGC-1 alpha (Addgene, 10974) using primers containing BamHI and EcoRI cloning sites (PPARGC1A_BamHI_F, PPARGC1A_EcoRI_R, see Supplementary Table 1) and sub-cloned into pENTR1A (Invitrogen, A10462). A 3’ 3x FLAG tag was then inserted by site directed mutagenesis using a Q5^®^ Site-Directed Mutagenesis Kit (NEB, E0554S) and the primers PPARGC1A_FLAG_SDM_F and PPARGC1A_FLAG_SDM_R (see Supplementary Table 1). PPARGC1A-3xFLAG was then cloned into pINDUCER20 (Addgene, 44012) using Gateway™ LR Clonase™ II Enzyme mix (Invitrogen, 11791-020), forming the vector pINDUCER20-PPARGC1A-3xFLAG.

shRNA vectors were created by annealing shRNA overlapping oligonucleotides and then cloned into pLKO.1 (Addgene, 10878). shRNA oligonucleotides sequences are detailed in Supplementary Table 1. Scramble shRNA pLKO.1 vector (Addgene, 1864) was used as a control.

Lentivirus was produced by as described previously (Tiscornia et al., 2006). Briefly, 3×10^6^ HEK293T cells were seeded in T75 flasks. The following day, HEK293T cells were transfected with 2.25 μg psPAX2 (Addgene, 12260), 1.5 pMD2.G (Addgene, 12259) and 3 μg target vector using PolyFect transfection reagent (Qiagen, 301107). Media containing virus was collected both 48 and 72 hours post-transfection and viral particles were precipitated using PEG-it™ Virus Precipitation Solution (System Biosciences, LV810A-1). To transduce cells with virus cells were treated with both virus (MOI 0.5-1.0) and 5 μg / mL Polybrene infection reaction (EMD Millipore, TR-1003). For pLKO.1 vectors, polyclonal cells were selected using 500 ng / mL puromycin (Sigma P7255) for 2 weeks; for pINDUCER20 vectors, polyclonal cells were selected using 250 μg / mL G418 (ThermoFisher Scientific, 10131027) for 2 weeks.

### Western blots

Cells were lysed in RIPA buffer (150 mM NaCl, 1% IGEPAL CA-630, 0.5% sodium deoxycholate, 0.1% SDS, 50 mM Tris pH 8.0, 1 mM EDTA) supplemented with protease inhibitors (Roche, 11836170001). Protein concentration was determined by BCA assay (Pierce, 23227). 5x SDS loading buffer (235 mM SDS, 50% glycerol, 0.005% bromophenol blue, 10% β-mercaptoethanol, 210 mM Tris-HCl pH 6.8) was then added to protein lysates to a 1x concentration and then incubated at 90°C for 10 minutes. Samples were then analysed by SDS-PAGE on 8% polyacrylamide gels using a PageRuler™ Prestained Protein Ladder (Thermo Scientific, 26616). Proteins where then transferred onto a nitrocellulose membrane (GE Healthcare, 10600002) and blocked using Odyssey^®^ Blocking Buffer (LI-COR Biosciences, P/N 927-40000). Primary antibodies used: anti-ERBB2 (Thermo Fisher, MA5-14057, 1:1,000), anti-phospho-ERBB2 (Sigma-Aldrich, SAB4300061, 1:1,000), anti-ERK (Cell Signaling Technologies, 4695S, 1:1,000), anti-phospho-ERK (Cell Signaling Technologies, 9106S, 1:2,000), anti-AKT (Cell Signaling Technologies, 2920S, 1:2,000), anti-phospho-AKT (Cell Signaling Technologies, 4060S, 1:2,000), anti-HNF4A (R&D Systems, PP-H1415-00, 1:1,000), anti-PPARGC1A (Novus Biologicals, NBP1-04676, 1:1,000), anti-phospho-PPARGC1A (Novus Biologicals, AF6650, 1:1,000), anti-GFP (Santa Cruz, sc-8334, 1:2,000). For the anti-PPARGC1A antibody, membranes were blocked and the primary antibody was diluted in 5% (w/v) milk powder in PBS. Secondary antibodies used: anti-rabbit (LI-COR Biosciences, 926-32213, 1:10,000) and anti-mouse (LI-COR Biosciences, 926-32210, 1:10,000). The membranes were visualized using a LI-COR Odyssey^®^ CLx Infrared Imager. Western blots were quantified using Empiria studio v1.1.

### RNA extraction and RT-qPCR

Total RNA was extracted from cells using a RNeasy Plus RNA extraction kit (Qiagen, 74136) according to the manufacturer’s protocol. RT-qPCR reactions were run using a QuantiTect SYBR Green RT-qPCR kit (Qiagen, 204243) on a Qiagen Rotorgene Q. Relative copy number of transcripts was determined from a standard curve and then normalised by the expression of *RPLP0* control gene. Primer pairs are listed in Supplementary Table 1.

### Seahorse metabolism assays

1×10^4^ OE19 cells were reverse transfected with siRNAs and seeded into 96-well plates (Agilent, 102601-100). 24 hours post-transfection cells were treated with 500 nM lapatinib or vehicle control. 24 hours after drug treatment normal culture medium (RPMI 1640) was changed to Seahorse XF RPMI assay medium (Agilent, 103681-100) and a mitochondrial stress test (Agilent, 103015-100) was carried out according to the manufacturer’s protocols using a Seahorse XFe96 analyser (Agilent) to measure oxygen consumption and extracellular acidification rates. Following completion of the mitochondrial stress test, a crystal violet assay was performed and oxygen consumption and extracellular acidification results were normalised by crystal violet 570 nm absorbance readings. Concentrations of inhibitors used: 1.5 μM oligomycin, 1 μM FCCP and 0.5 μM rotenone/antimycin A.

### ATAC-seq processing and analysis

Two biological replicates were sequenced per condition. Omni-ATAC-seq was performed following published protocols (Corces et al., 2017). Cells were dissociated from plates using trypsin (Gibco, 25300-054) and 2×10^5^ cells were collected and centrifuged at 500 RCF for 5 minutes. Cells were then washed with PBS and centrifugation repeated. The omni-ATAC-seq protocol (Corces et al., 2017) was then followed until the isolation of nuclei. Nuclei were centrifuged at 500 RCF, 4 °C for 10 minutes and re-suspended in 10 μL nuclease free water (Ambion, AM9937). Nuclei were then counted using a haemocytometer, and 5×10^4^ nuclei were used for the transposition reaction. Libraries were size selected using Ampure XP beads (Beckman Coulter Agencourt, A63881) using a two-sided selection (0.5x reaction volume and 1.25x reacton volume) and eluted in 12 μL nuclease free water. ATAC-seq libraries were then sequenced by the University of Manchester Genomic Technologies Core Facility on a HiSeq 4000 System (Illumina).

Initial processing of ATAC-seq was performed as described previously (Britton et al., 2017). Reads were trimmed using Trimmomatic v0.32 (Bolger et al., 2014) and aligned to the human genome (GRCh37, hg19) using Bowtie2 v2.3.0 (Langmead and Salzberg, 2012) with the following options: -X 2000 -dovetail. Using SAMtools v1.9 (Li et al., 2009), only mapped reads(>q30) were retained. Reads mapping to blacklisted regions were removed using BEDtools v2.27.1 (Quinlan and Hall, 2010). Duplicates were then marked using Picard (https://broadinstitute.github.io/picard/). Peaks were called using MACS2 v2.1.1 (Zhang et al., 2008) with the following parameters: -q 0.01, -nomodel-shift −75 -extsize 150 -B -SPMR. For the drug withdrawal timecourse all samples were processed as described above but the - SPMR option was not used when calling peaks with MACS2.

To create a union peakset for each experiment, biological replicates were checked for concordance (r > 0.90) and then merged into a single alignment file. Peaks were then re-called using MACS2 and the top 50,000 most significant peaks from each condition were retained. Peak summits were extended +/− 250 bp using BEDtools slop and then a union peakset of all conditions was created using HOMER v4.9 (Heinz et al., 2010) mergePeaks.pl using the -d 250 parameter. This union peakset was then used to identify differentially accessible regions by counting reads mapping to peaks (i.e. accessible regions) in individual replicates using featureCounts v1.6.2 (Liao et al., 2014). Read counts were then used in DESeq2 v1.14.1 (Love et al., 2014) to call differentially accessible regions. Typically, an FDR (adjusted p-value) < 0.05 and 2 fold linear fold change cut off was used to define differential regions. To identify transcription factor binding motifs enriched in differentially accessible regions, peaks were separated into promoter or non-promoter associated peaks based on whether the peaks were −2.5 kb / + 0.5 kb from the TSS using BEDtools intersectBed. De novo or known motif enrichment was then performed in non-promoter peaks using HOMER v4.9 (Heinz et al., 2010).

### ChIP-qPCR and ChIP-seq processing and analysis

ChIP-qPCR and ChIP-seq was performed as described previously (Rogerson et al., 2020). Primers for ChIP-qPCR are listed in Supplementary Table 5. 5 × 10^6^ nuclei and 2.5 μg antibody were used for transcription factor/co-activator immunoprecipitation, and 2 × 10^6^ nuclei and 2 μg antibody were used for histone marks. 50 μL of protein A or G Dynabeads were used for immunoprecipitation (Invitrogen, 10002D and 10004D). Antibodies used: anti-H3K27ac (abcam, ab4729), anti-HNF4A (R&D Systems, PP-H1415-00), anti-PPARGC1A (Novus Biologicals, NBP1-04676). For ChIP-seq, two biological replicates were sequenced per condition and replicates were checked for concordance (r > 0.80). Whilst spike in control chromatin was supplemented to chromatin, analysis of results showed that ‘reads in peaks’ normalisation of HNF4A and H3K27ac ChIP-seq data was more appropriate because global changes to the levels of HNF4A and H3K27ac were not observed.

Reads were trimmed using Trimmomatic v0.32 (Bolger et al., 2014) and aligned to the human genome (GRCh37, hg19) using Bowtie2 v2.3.0 (Langmead and Salzberg, 2012). Mapped reads (>q30) were retained using SAMtools v1.9 (Li et al., 2009). Reads mapping to blacklisted regions were removed using BEDtools v2.27.1 (Quinlan and Hall, 2010). Duplicates were then marked using Picard (https://broadinstitute.github.io/picard/). Peaks were called using MACS2 v2.1.1, using input DNA as control (Zhang et al., 2008). For differential binding analysis peak summits were extended +/− 250 bp using BEDtools slop and a union peakset was created by merging peaks from each sample using HOMER mergePeaks.pl (-d 250). Reads mapping to peaks in the union peakset were then counted using featureCounts v1.6.2 and analysed in DESeq2 v1.14.1 to call differentially bound sites (FDR < 0.1).

### RNA-seq processing and analysis

Three biological replicates were sequenced per condition. Total RNA was extracted from cells using a RNeasy Plus RNA extraction kit (Qiagen, 74136). An on-column DNase digest (Qiagen, 79254) was performed according to the manufacturer’s protocol. RNA-seq libraries were generated using a TruSeq stranded mRNA library kit (Illumina, RS-122-2001) and sequenced by the University of Manchester Genomic Technologies Core Facility on a HiSeq 4000 System (Illumina).

Reads were trimmed using Trimmomatic v0.32 (Bolger et al., 2014) and aligned to the human genome (GRCh37, hg19, RefSeq transcript annotation) using STAR v2.3.0 (Dobin et al., 2012). Gene expression counts were obtained using featureCounts v1.6.2 (Liao et al., 2014) and differentially expressed genes were identified using DESeq2 v1.14.1 using FDR < 0.05 (Love et al., 2014). For results from the lapatinib treatment timecourse, differentially expressed genes were filtered to remove genes in which no timepoint had an FPKM value > 1. Metascape (Zhou et al., 2019) was used for gene ontology analysis of differentially expressed genes. Ingenuity Pathway Analysis (Krämer et al., 2014) was used to predict upstream regulators.

Patient tumour samples in the OCCAMS dataset expressing *ERBB2* at high levels (*ERBB2*^HIGH^) were defined by *ERBB2* expression levels greater than the median +2 SD.

### Bioinformatics and data visualisation

To visualise data, the bigwig files from an experiment were normalised using the reciprocal of scale factors obtained in DESeq2. The bigwigCompare tool from deepTools v3.1.1 (Ramírez et al., 2016) was then used to scale individual replicate bigwigs. Bigwigs for each condition were then merged using UCSC bigwigmerge and the resulting bedgraph files were converted to bigwigs for visualisation in IGV v2.7.2 (Robinson et al., 2011). Heatmaps of epigenomic data were generated using deepTools. Tag density plots were generated in deepTools and the data was then plotted in Microsoft Excel. Heatmaps of expression data and Pearson correlations were generated using Morpheus (https://software.broadinstitute.org/morpheus/). Peaks were annotated to genes using HOMER for the nearest gene model or GREAT (McLean et al., 2010) for the basal plus extension model. Principal component analysis was performing using the prcomp function in R v3.6.0. Euler diagrams were generated using the R Eulerr library.

### Statistical tests

To determine whether an overlap of genes or peaks is statistically significant a hypergeometric test was used, using the dhyper function in R v3.6.0. Other statistical tests were performed using GraphPad Prism v8. All T-tests were two-tailed.

### Datasets

All data was obtained from ArrayExpress, unless stated otherwise. Cell line RNA-seq data was obtained from: EBI, EGAD00001001357 (Cancer Genome Project, cancer.sanger.ac.uk, Tate et al., 2019); NCI-N87 RNA-seq data, Sequence Read Archive SRP091839 (Zeng et al., 2018); OE33 RNA-seq data, E-MTAB-5175 (Britton et al., 2017). HET1A ATAC-seq data, E-MTAB-6931 (Rogerson et al., 2019). CP-A ATAC-seq data, E-MTAB-8994 (Rogerson et al., 2020).

Patient tissue RNA-seq data was obtained from: E-MTAB-4054 (Maag et al., 2017) and the OCCAMS consortium (EGAD00001007496). Human tissue ATAC-seq data was obtained from: E-MTAB-5169 (Britton et al., 2017), E-MTAB-6751 (Rogerson et al., 2019), E-MTAB-8447 (Rogerson et al., 2020) and The Cancer Genome Atlas OAC ATAC-seq data were obtained from the GDC data portal (portal.gdc.cancer.gov; Corces et al., 2018).

OE19 HNF4A ChIP-seq data was obtained from E-MTAB-6858 (Rogerson et al., 2019). OE19 siERBB2 RNA-seq data was obtained from E-MTAB-8579 (Rogerson et al., 2020). OE19 H3K27ac ChIP-seq data was obtained from NCBI SRA SRP201335 (Chen et al., 2020). NRF1 ChIP-seq data was obtained from ENCODE: HepG2, ENCSR853ADA; K562, ENCSR837EYC; MCF7, ENCSR135ANT.

### Data access

Sequencing data have been deposited in ArrayExpress. OE19 lapatinib treatment timecourse ATAC- and RNA-seq: E-MTAB-10302, E-MTAB-10304. Lapatinib treatment of WTSI-OESO_009, ESO26, KYAE1 and NCI-N87 cells ATAC-seq: E-MTAB-10306, E-MTAB-10307, E-MTAB-310, E-MTAB-313. Lapatinib withdrawal timecourse ATAC-seq: E-MTAB-10314. OE19 siPPARGC1A RNA-seq: E-MTAB-10317. OE19 HNF4A, PPARGC1A and H3K27ac ChIP-seq data: E-MTAB-10334.

## Acknowledgements

We thank Guanhua Yan for excellent technical assistance, Connor Rogerson for advice and help with RNA-seq data processing and staff in the Bioinformatics, Genomic Technologies and Flow cytometry core facilities. Rebecca Fitzgerald for providing the OAC organoid. We also thank Nicoletta Bobola, Tim Somervaille and Shen-Hsi Yang for critical appraisal of the manuscript. This work was funded by grants from the Wellcome Trust (103857/Z/14/Z and 102171/Z/13/Z).

Author contributions: S.O. and K.C. performed the experiments and data analysis in this study; J.B. led the metabolic studies; A.D.S. led the experimental parts of the program. All authors contributed to manuscript preparation and/or critically appraised manuscript drafts.

