## Supplementary Figures for "Widespread reorganisation of the regulatory chromatin landscape facilitates resistance to inhibition of oncogenic ERBB2 signalling"

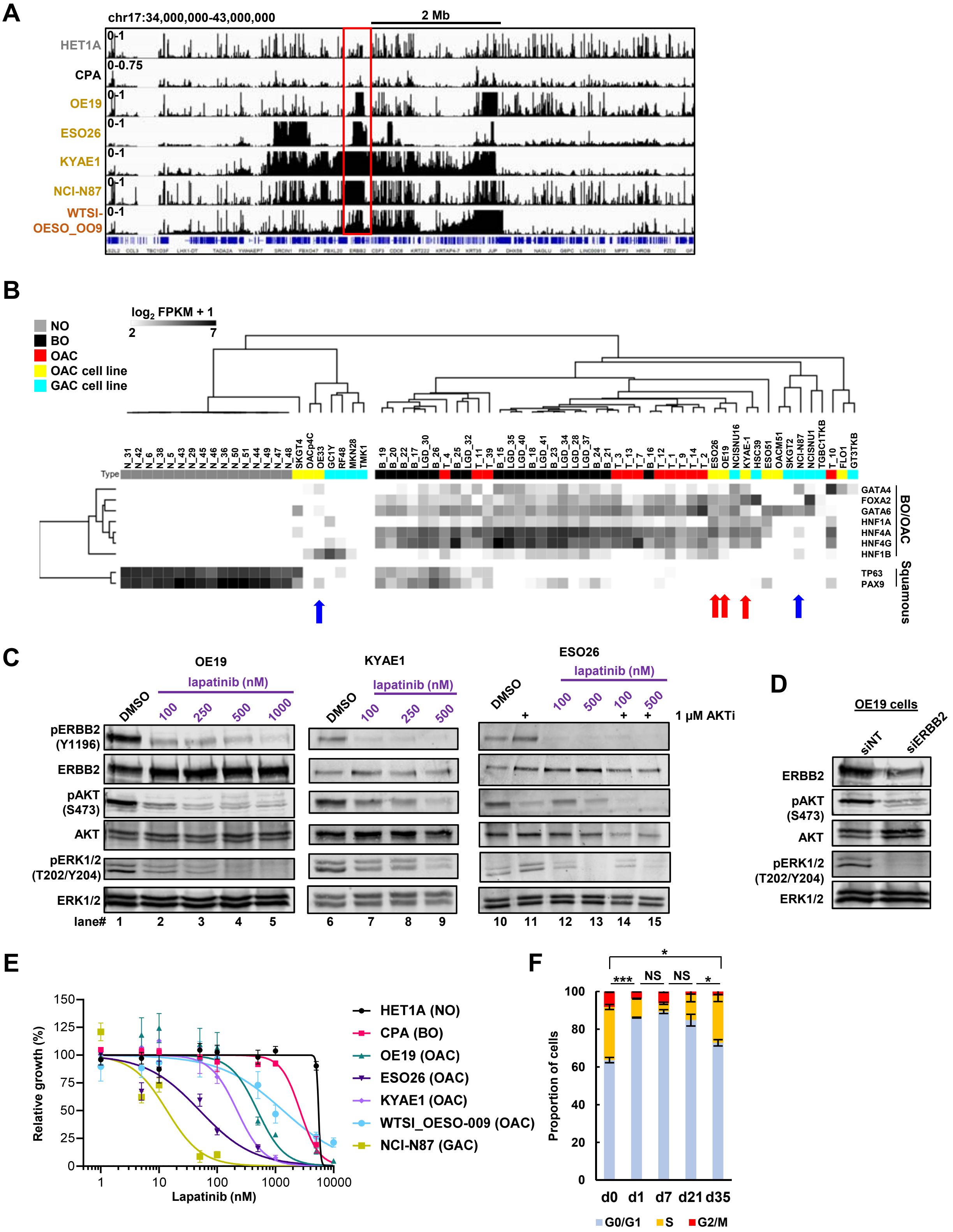

**Figure S1. Resistance to ERBB2 inhibition in OAC cells arises after weeks of treatment.** (A) Genome browser view of ATAC-seq data highlighting the *ERBB2* locus in the indicated cell lines and organoid. (B) Heatmap showing expression of selected genes in patient tissue, OAC cell lines and gastric adenocarcinoma (GAC) cell lines. Arrows represent cell lines investigated in this study. (C) Western blot of OE19, KYAE1 and ESO26 cells treated with DMSO or the indicated doses of lapatinib for 24 hours. ESO26 cells were also treated with 1 μM AKT inhibitor (AKTi – MK-2206) as indicated. (D) Western blot of OE19 cells treated with ERBB2 siRNA (siERBB2) or non-targeting control (siNT) for 72 hours. (E) MTS growth assay of cell lines treated with lapatinib for 72 hours. NO – normal oesophageal, BO – Barrett's oesophageal, OAC – oesophageal adenocarcinoma, GAC – gastric adenocarcinoma. (F) Cell cycle analysis of OE19 cells treated with vehicle control (d0) or 500 nM lapatinib for the indicated number of days. Statistical significance was determined using an unpaired T-test on the proportion of cells in GO/G1; *** $P < 0.001$, * $P < 0.05$, NS – not significant, $n = 3$, error bars depict SEM.

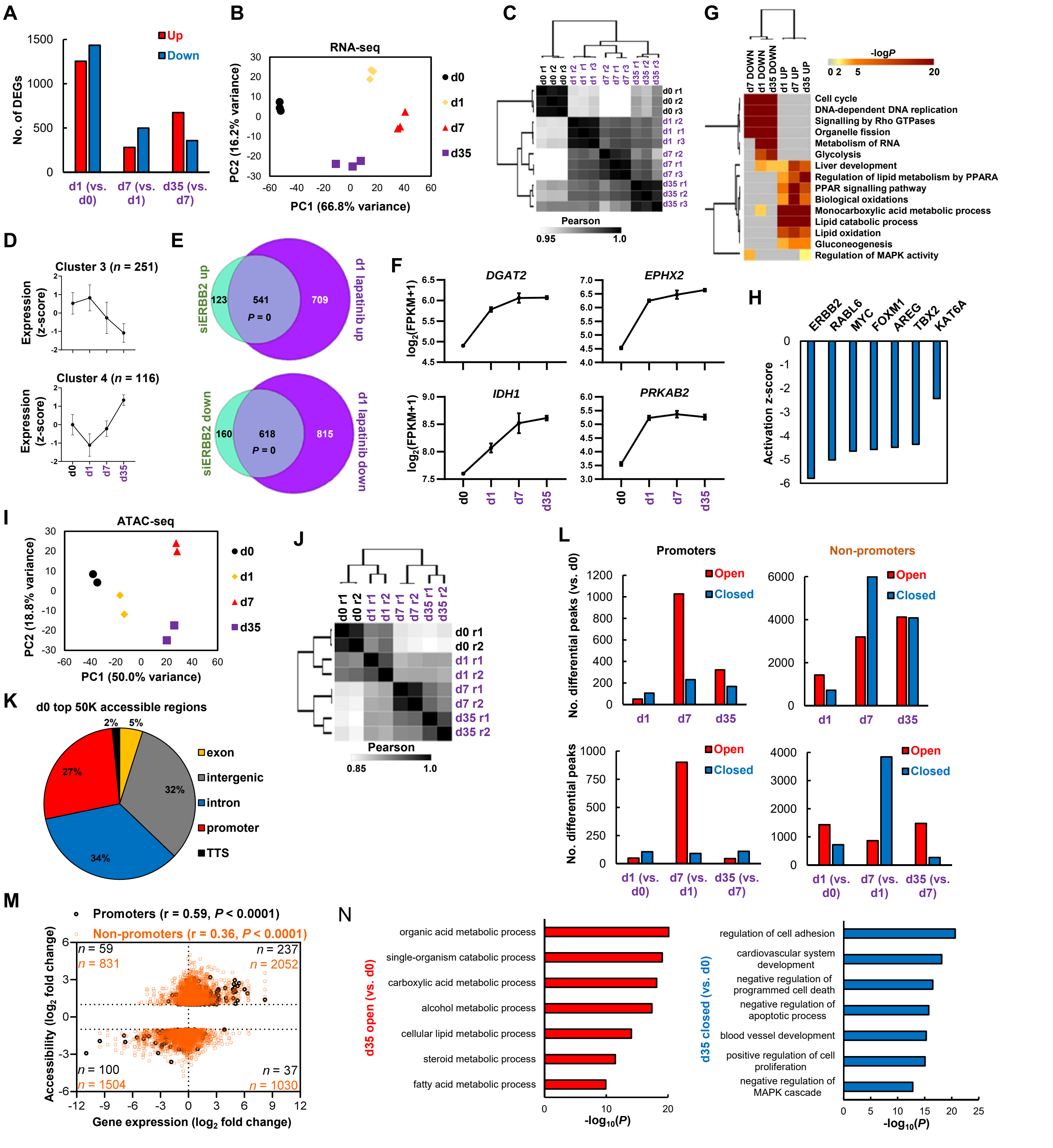

**Figure S2. ERBB2 inhibition causes global changes to the transcriptome and chromatin landscape of OAC cells.**
(A) The number of DEGs in OE19 cells after 500 nM lapatinib treatment, relative to the earlier timepoint. (B) Principal component analysis and (C) Pearson correlation analysis of OE19 RNA-seq data for samples treated with 500 nM lapatinib for the indicated number of days (d0-d35) and replicates (r1-3). (D) Average expression profiles of genes in the indicated clusters from Fig. 2C across the lapatinib treatment time course. Error bars depict SD. (E) Overlap of DEGs (2X linear fold change, FDR < 0.05, FPKM > 1) in OE19 cells after 72 hours siERBB2 treatment or 24 hours 500 nM lapatinib treatment. Statistical significance was determined using a hypergeometric test. (F) Expression of example up-regulated genes across the lapatinib treatment timecourse. (G) GO analysis using Metascape of DEGs in OE19 cells after lapatinib treatment for the indicated timepoints relative to control (d0) cells. (H) Ingenuity pathway analysis predicting inhibited upstream regulators in resistant OE19 cells (d35) relative to control (d0) cells. (I) Principal component analysis and (J) Pearson correlation analysis of OE19 ATAC-seq data. (K) Genomic distribution of top 50,000 ATAC-seq peaks (i.e. accessible chromatin regions) in control (d0) OE19 cells. (L) The number of differentially accessible peaks in promoter or non-promoter regions either relative to control cells (d0) (top panel) or the previous timepoint (bottom panel). (M) Comparison of the fold change of differentially accessible peaks (2X linear fold change, FDR < 0.05) and the fold change of genes in resistant OE19 cells (d35) relative to control (d0) cells. Differential peaks were annotated to genes using the nearest gene model. Peaks are partitioned into promoter (black) or non-promoter (orange) regions. The number of peaks ($n$) in each quadrant is indicated. Pearson correlation (r) is shown. (N) GO analysis (GREAT) of genes annotated to d35 (vs. d0) differentially accessible peaks using the basal plus extension model.

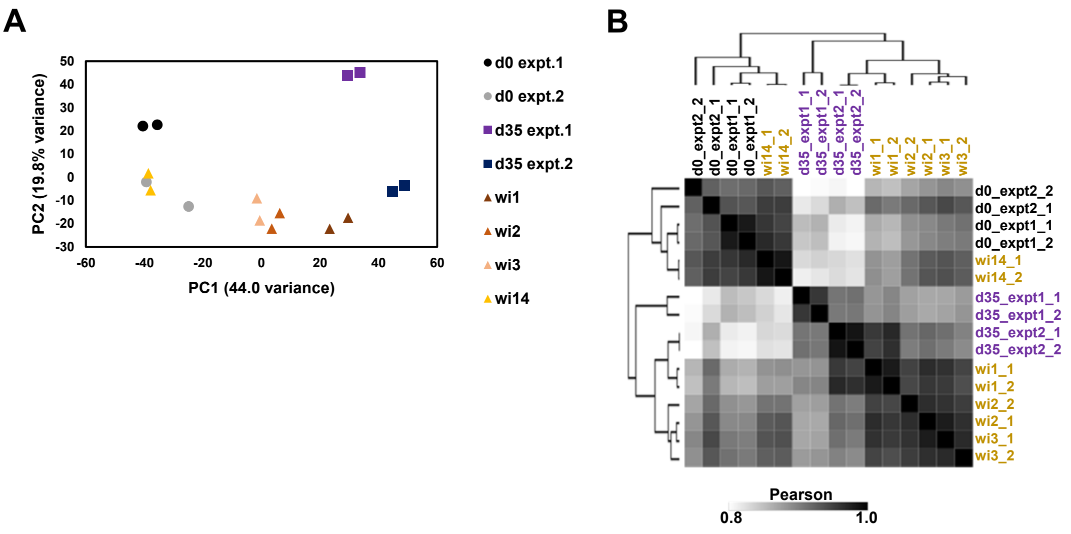


**Figure S3. Chromatin accessibility changes in resistant cells are reversible following drug withdrawal.**
(A and B) Heatmap showing Pearson correlation (A) and principal component analysis (B) of OE19 ATAC-seq data from both the lapatinib treatment and drug withdrawal timecourses.

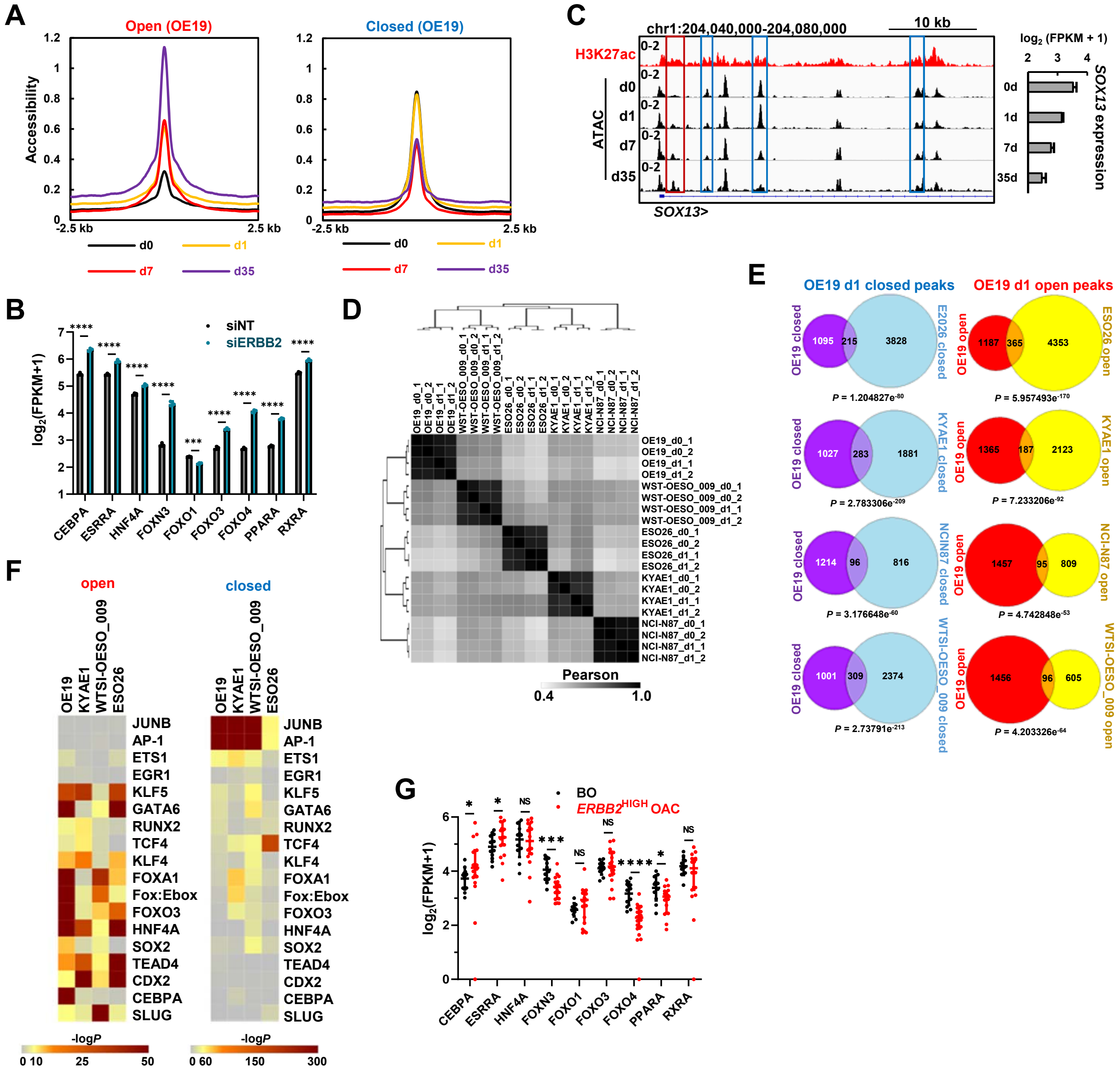

**Figure S4. A gastro-intestinal transcription factor programme is activated after ERBB2 inhibition in OAC cells.** (A) Tag density plot of ATAC-seq signal centred on differentially open or closed chromatin regions after 35 days lapatinib treatment (d35) relative to control (d0) cells. (B) Expression of the indicated transcription factors in OE19 cells treated with ERBB2 siRNA (siERBB2) or non-targeting control (siNT) for 72 hours. (C) Genome browser view of the *SOX13* locus in OE19 cells showing ATAC-seq and H3K27ac ChIP-seq (Chen et al., 2020) data. Differentially open (red) or closed (blue) are highlighted using rectangles. Bar chart depicts *SOX13* expression over the lapatinib treatment timecourse. (D) Pearson correlation of ATAC-seq data of gastro-oesophageal cells treated with DMSO (d0) or lapatinib (d1) for 1 day. Doses of lapatinib used: OE19, ESO26 and KYAE1 – 500 nM, WTSI_OESO-009 – 1 μM, NCI-N87 250 nM. ESO26 (*PIK3CA* activating mutation) were treated with lapatinib and 1 μM MK-2206 (AKTi). (E) Overlap of differentially open or closed peaks in OE19 cells and indicated cell lines. Statistical significance was determined using a hypergeometric test. (F) Heatmap of *P*-values of known motifs enriched in differentially open or closed chromatin after lapatinib treatment (d1) relative to control (d0) cells in the indicated OAC cells. (G) Expression of the indicated transcription factors in BO and *ERBB2*$^{HIGH}$ OAC tissue.

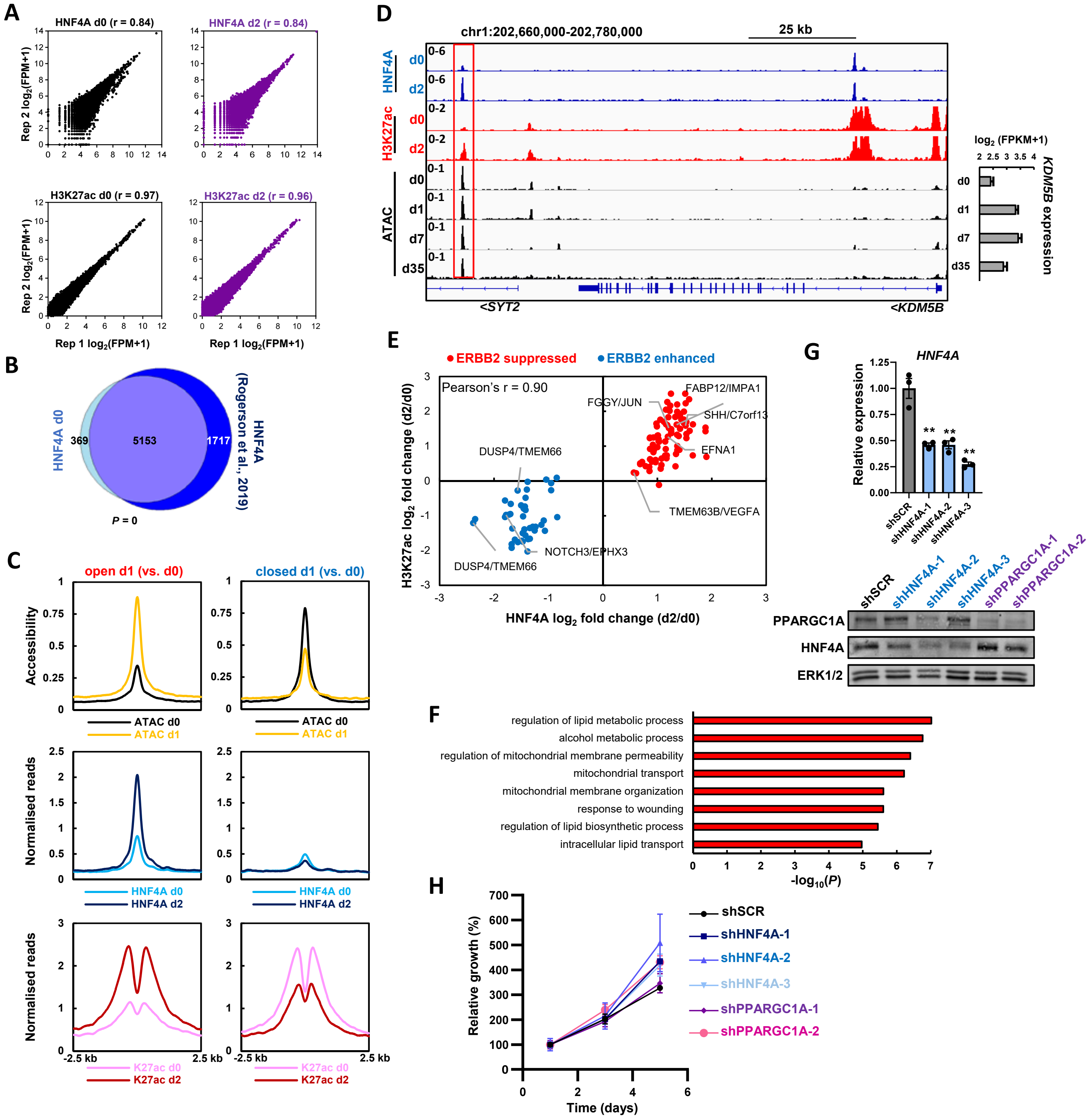


**Figure S5. HNF4A regulatory activity is activated/altered following ERBB2 inhibition?**
(A) Correlation of biological replicates from HNF4A and H3K27ac ChIP-seq in OE19 cells treated with DMSO (d0) or 500 nM lapatinib (d2) for 2 days. (B) Overlap of HNF4A binding sites in OE19 cells treated with DMSO for 2 days (d0) and published OE19 HNF4A ChIP-seq data (Rogerson et al., 2019). Biological replicates were intersected and then compared. Statistical significance was determined using a hypergeometric test. (C) Tag density plot of ATAC-seq, HNF4A binding and H3K27ac signal at differentially open or closed peaks after 1 day lapatinib treatment (d1) relative to control cells (d0). (D) Genome browser view of the *KDM5B* locus in OE19 cells. Red box highlights a putative enhancer in which HNF4A binding increases after 2 days lapatinib treatment. *KDM5B* mRNA expression over the lapatinib timecourse is shown on the right. (E) Scatter plot of $\log_2$ fold change in HNF4A binding and H3K27ac signal after 2 days 500 nM lapatinib treatment. H3K27ac peaks were assigned to HNF4A peaks by extending HNF4A peaks by 500 bp in each direction and then intersecting with H3K27ac peaks. (F) GO term analysis of genes associated with HNF4A peaks (found in d2 lapatinib treated cells) located in lapatinib induced accessible peaks (d1 treatment). (G) RT-qPCR and western blot for HNF4A mRNA and protein levels in OE19 cells treated with the indicated shRNAs. ** $P < 0.01$, unpaired T-test, $n = 3$. (H) Crystal violet growth assay of OE19 cells treated with the indicated shRNAs.

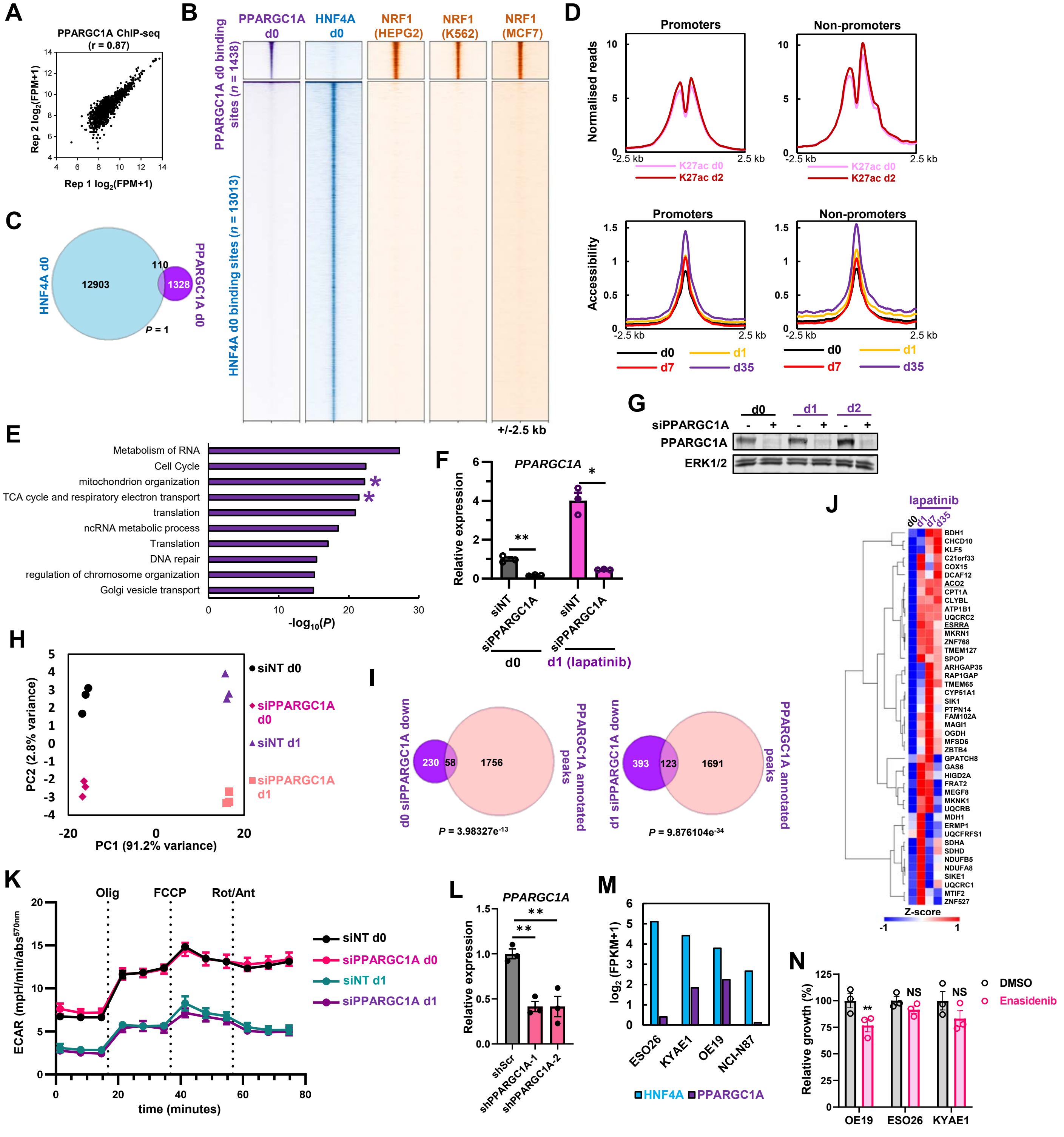


**Figure S6. PPARGC1A regulates mitochondrial processes and supports the emergence of resistance.**

(A) Scatter plot of $\log_2$(FPM+1) of biological replicates from OE19 PPARGC1A ChIP-seq in OE19 cells. (B) Heatmap of PPARGC1A and HNF4A binding sites in parental OE19 cells. NRF1 ChIP-seq signal in HEPG2, K562 and MCF7 cells is shown. (C) Overlap of HNF4A and PPARGC1A binding sites in parental OE19 cells. Statistical significance was determined using a hypergeometric test. (D) Tag density plots of H3K27ac signal (top) and ATAC-seq signal (bottom) at PPARGC1A binding sites in promoters or non-promoter regions. (E) GO analysis (Metascape) of genes annotated to PPARGC1A binding sites using the basal plus extension model (GREAT). (F) RT-qPCR analysis of OE19 cells treated with non-targeting control (siNT) or siPPARGC1A (siPP). Cells were also treated with DMSO (d0) or 500 nM lapatinib (d1) for 24 hours. ** $P < 0.01$, * $P < 0.05$, paired T-test, $n = 3$, error bars depict SEM. (G) Western blot analysis of OE19 cells treated with siNT or siPPARGC1A and DMSO for 24 hours (d0) or 500 nM lapatinib for 24 (d1) or 48 (d2) hours. (H) Principal component analysis of RNA-seq data from OE19 cells treated with siPPARGC1A for 2 days and also treated with DMSO (d0) or 500 nM lapatinib (d1) for 1 days. (I) Overlap of siPPARGC1A down-regulated genes (FDR < 0.05) in the presence (d1) or absence (d0) of lapatinib and genes annotated to PPARGC1A binding sites. Overlapping genes are PPARGC1A direct target genes. Statistical significance was determined using a hypergeometric test. (J) Heatmap of RNA-seq data of PPARGC1A direct target genes up-regulated by lapatinib (FDR < 0.05, up-regulated d1 or d7 or d35 relative to d0, $n = 45$) in OE19 cells. (K) OE19 glycolytic rate during a mitochondrial stress test. Cells were treated with siNT or siPPARGC1A, and DMSO or 500 nM lapatinib for 24 hours. ECAR – extracellular acidification rate. (L) RT-qPCR for *PPARGC1A* mRNA expression in OE19 cells treated with the indicated shRNAs, ** $P < 0.01$, unpaired T-test, $n = 3$. (M) Expression from RNA-seq data of *HNF4A* and *PPARGC1A* in the indicated cell lines. (N) Crystal violet growth assay of the indicated cell lines after treatment with DMSO or 5 μM enasidenib for 72 hours. ** $P < 0.01$, paired T-test, $n = 3$, error bars depict SEM.
